# Diagnostic high-throughput sequencing of 2,390 patients with bleeding, thrombotic and platelet disorders

**DOI:** 10.1101/504142

**Authors:** Kate Downes, Karyn Megy, Daniel Duarte, Minka Vries, Johanna Gebhart, Stefanie Hofer, Olga Shamardina, Sri VV Deevi, Jonathan Stephens, Rutendo Mapeta, Salih Tuna, Namir Al Hasso, Martin W Besser, Nichola Cooper, Louise Daugherty, Nick Gleadall, Daniel Greene, Matthias Haimel, Howard Martin, Sofia Papadia, Shoshana Revel-Vilk, Suthesh Sivapalaratnam, Emily Symington, Will Thomas, Chantal Thys, Alexander Tolios, Christopher J Penkett, NIHR BioResource, Willem H Ouwehand, Stephen Abbs, Michael A Laffan, Ernest Turro, Ilenia Simeoni, Andrew D Mumford, Yvonne MC Henskens, Ingrid Pabinger, Keith Gomez, Kathleen Freson

## Abstract

A targeted high-throughput sequencing (HTS) panel test for clinical diagnostics requires careful consideration of the inclusion of appropriate diagnostic-grade genes, the ability to detect multiple types of genomic variation with high levels of analytic sensitivity and reproducibility, and variant interpretation by a multi-disciplinary team (MDT) in the context of the clinical phenotype. We have sequenced 2,390 index patients using the ThromboGenomics HTS panel test of diagnostic-grade genes known to harbour variants associated with rare bleeding, thrombotic or platelet disorders (BPD). The diagnostic rate was determined by the clinical phenotype, with an overall rate of 50.4% for all thrombotic, coagulation, platelet count and function disorder patients and a rate of 6.2% for patients with unexplained bleeding disorders characterized by normal hemostasis test results. The MDT classified 756 unique variants, including copy number and intronic variants, as Pathogenic, Likely Pathogenic or Variants of Uncertain Significance. Almost half (49.7%) of these variants are novel and 41 unique variants were identified in 7 genes recently found to be implicated in BPD. Inspection of canonical hemostasis pathways identified 29 patients with evidence of oligogenic inheritance. A molecular diagnosis has been reported for 897 index patients providing evidence that introducing a HTS genetic test for BPD patients is meeting an important unmet clinical need.

**Key points:**

1. High-throughput sequencing (HTS) test reveals a molecular diagnosis for 38% of 2,390 patients with bleeding, thrombotic and platelet disorders.
2. ThromboGenomics HTS test validates recent gene discoveries and detects copy number and intronic variants.

## Introduction

Inherited bleeding, thrombotic and platelet disorders (BPD) are a heterogeneous group of rare disorders caused by DNA variants in a large number of loci. The most common bleeding disorders are von Willebrand Disease (VWD), affecting up to 0.01% of the population and Hemophilia A and B, which together affect 0.01% of males.^1^ There are no accurate estimates of the prevalence of the remaining rare inherited bleeding disorders, although registry data suggest the prevalence being <0.001%.^2^ Venous thrombosis has an overall annual incidence of less than 1 in 1,000, however it is rare in the pediatric population, with rates of approximately 1 in 100,000, indicative of possible environmental and lifestyle effects in adult patients.^3^ To obtain a conclusive molecular diagnosis requires attendance at multiple outpatient consultations for a large portion of patients with an assumed diagnosis of a rare inherited BPD.

The genetic architecture of inherited BPDs is well determined, however, new genes continue to be identified. To date there are 21, 11 and 63 diagnostic-grade genes (hereafter TIER1 genes) associated with coagulation, thrombotic and platelet disorders, respectively.

Since validation of the ThromboGenomics HTS test^4^, 33 TIER1 genes, including recently discovered BPD genes, have been added to the HTS test, increasing the clinical utility. Others have reported on similar gene panel tests or used whole exome sequencing to identify DNA variants causing inherited BPDs^5–9^. However, all these studies have reported on relatively small numbers of patients (<160 index patients) preventing firm conclusions about the clinical utility of such tests. The diagnostic rates obtained in these studies cannot be compared as all focused on different sets of genes (with a subset of TIER1 genes, and also inclusion of ‘research’ genes), used different patient inclusion criteria and variant classification was not standardised as is now recommended.^10^

Here we report on the results obtained with the ThromboGenomics HTS test for 2,390 index patients categorised into five classes of disorders based on the appended human phenotype ontology (HPO) terms; ^11^ thrombotic, platelet count, platelet function, coagulation and unexplained bleeding. Variant interpretation, by a multi-disciplinary team (MDT), determined the contribution of variants in TIER1 genes to the observed phenotypes thereby providing insights in the clinical utility of HTS testing for different categories of patients. We also comment on the standardisation of variant interpretation and how the reporting of a conclusive molecular diagnosis has immediately impacted on clinical management. Finally, due to the large number of patients tested, we are able to highlight the clinical importance of detecting copy number and deep intronic variants and possible oligogenic inheritance.

## Methods

### Patients

The 2,390 index patients were either referrals for the ThromboGenomics test or patients who joined the PANE and VIBB studies (Supplemental Table 1). Clinical and laboratory phenotypes were recorded using HPO terms as described.^4,12^ Further details of the study participants, including institutional review board or research ethics committee information are in the Supplemental Information.

#### ThromboGenomics referrals for diagnostic testing of inherited BPDs

Samples and clinical/laboratory phenotype information from 1,602 index patients with a known or suspected diagnosis of inherited BPD according to criteria as described were referred by clinicians in 72 UK and 46 non-UK hospitals (Supplemental Figure 1).^4^

#### PANE: Preoperative screening for mild bleeding risk

A total of 212 patients, identified through preoperative assessment of bleeding risk at Maastricht University Medical Centre (MUMC), were recruited and underwent a full hematological assessment including extensive laboratory testing for hemostasis parameters (Supplemental Table 2).^13,14^

#### VIBB: Vienna Bleeding Biobank

A total of 594 patients referred to the Hematology and Hemostaseology specialist tertiary referral centre in Vienna for assessment of a mild to moderate bleeding disorder were recruited and subjected to a full hematological assessment, including extensive laboratory testing for hemostasis parameters (Supplemental Table 2).^15^

### ThromboGenomics HTS test

The ThromboGenomics HTS test sample preparation and sequencing protocols are as described with minor modifications (Supplemental Information).^4^ The content of the test has been reversioned twice since first described to include additional (mostly recently discovered) TIER1 genes. ThromboGenomics version 2 (TG.V2) and version 3 (TG.V3) include 80 and 96 genes, respectively (Supplemental Table 3). TIER1 genes are curated and approved by the Scientific and Standardization Committee on Genomics in Thrombosis and Hemostasis (SSC-GinTH) of the International Society on Thrombosis and Haemostasis (ISTH). For TG.V3, probes for 10,000 common single nucleotide variants (SNVs) were included to estimate relatedness and ancestry. Automated bioinformatics analysis pipeline methods, including variant calling, are described in the Supplemental Information.

### Variant prioritisation and interpretation

Variants were annotated and prioritised for interpretation using the analytical process as reported^4^ based on the predicted effect in the curated transcript, presence in the Human Gene Mutation Database (HGMD)^16^ or in a curated set of known pathogenic variants (Supplemental Information) and the minor allele frequency (MAF) in the Exome Aggregation Consortium (ExAC) and Genome Aggregation (gnomAD) databases.^17^ On a patient-by-patient basis, DNA variants passing filtering were prioritised and interpreted by a MDT in the context of the HPO terms and family history. Reported variants were characterised as Pathogenic, Likely Pathogenic and Variants of Uncertain Significance (VUS) alongside a decision of the likely contribution of each variant to the patients phenotype. The MDT made use of Congenica’s diagnostic decision support platform Sapientia™ (Cambridge, UK) to support the review process and record findings in the form of research reports for return to referring clinicians. For all samples sequenced using TG.V2, variant interpretation was performed according to guidelines agreed by the members of the ThromboGenomics MDT (criteria in Supplemental Table 4). In 2017, the UK Association of Clinical Genomic Science published best practice guidelines for variant interpretation based on the earlier reported American College of Medical Genetics and Genomics (ACMG) guidelines.^10^ Sapienta software implemented ACMG guidelines from release 1.7 (January 2018), allowing rapid variant interpretation following the guidelines. This updated system was applied for variant interpretation of all samples sequenced using TG.V3. Supplemental Table 5 summarises the main differences in panel content, methods, analysis and interpretation used for TG.V2 and TG.V3.

## Results

### Patient inclusion criteria and phenotypes

A total of 2,390 index patients and 156 samples from relatives and carriers were tested using the ThromboGenomics HTS test (Supplemental Table 1, Supplemental Figure 1). The largest group of 1,602 index patients and 18 referrals for hemophilia carrier status, were referred by specialist tertiary centres for diagnostic testing. To better appreciate the clinical utility of the HTS test we also included 193 and 594 samples from the PANE and VIBB single-centre studies. Both studies excluded patients with platelet counts <100×10^9^/l.

Based on the clinical and laboratory phenotypes of all patients, a total of 7,340 HPO terms were appended, which were used to categorise all patients into five broad disorder classes: thrombotic (n=285), platelet count (n=329), platelet function (n=397), coagulation (n=685) and unexplained bleeding (n=698) (Figure 1A). The ThromboGenomics referrals included patients of all five classes, while the majority of the PANE (80.3%) and VIBB (60.1%) patients are classed as unexplained bleeding.

**Figure 1.**
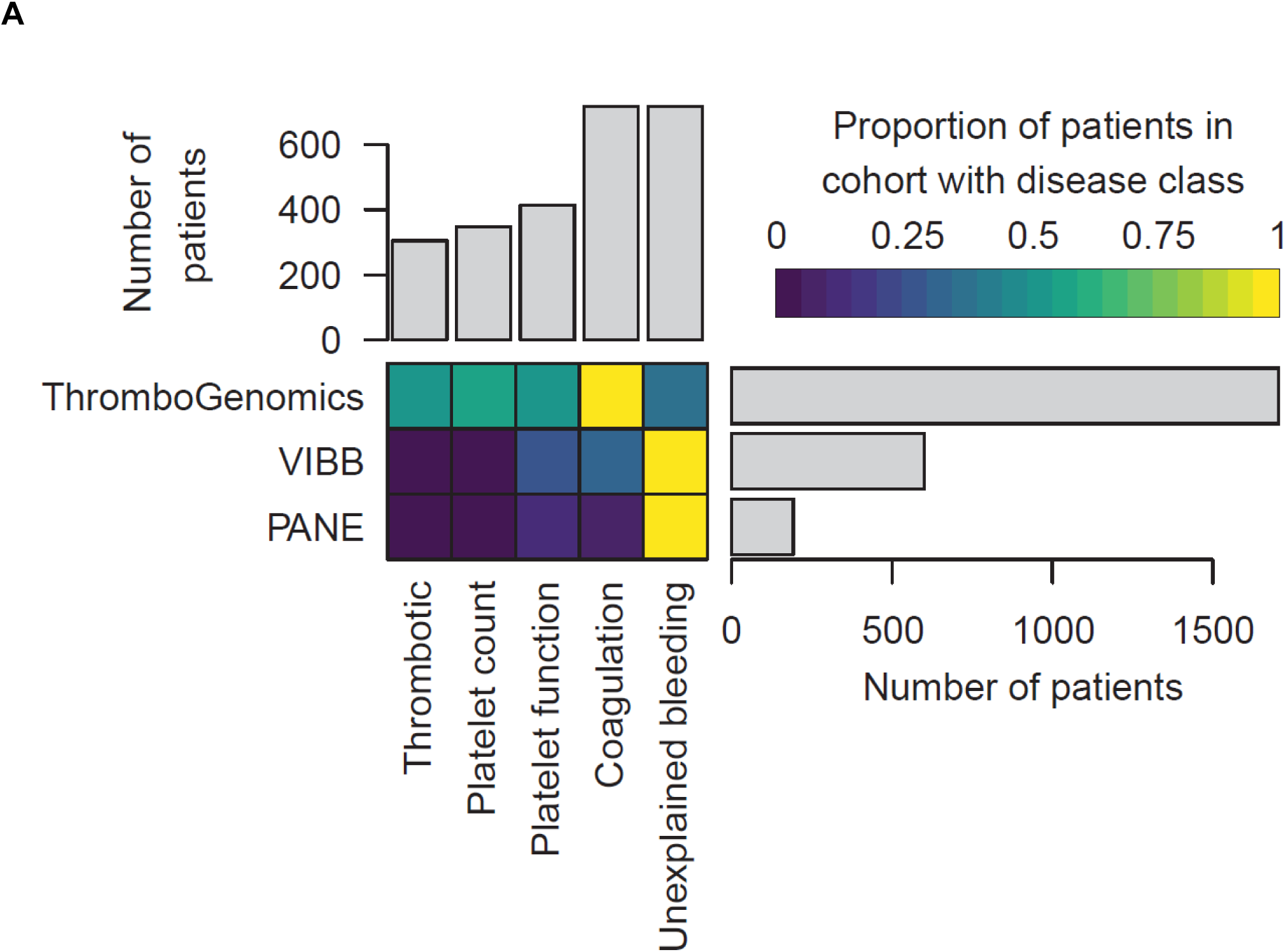
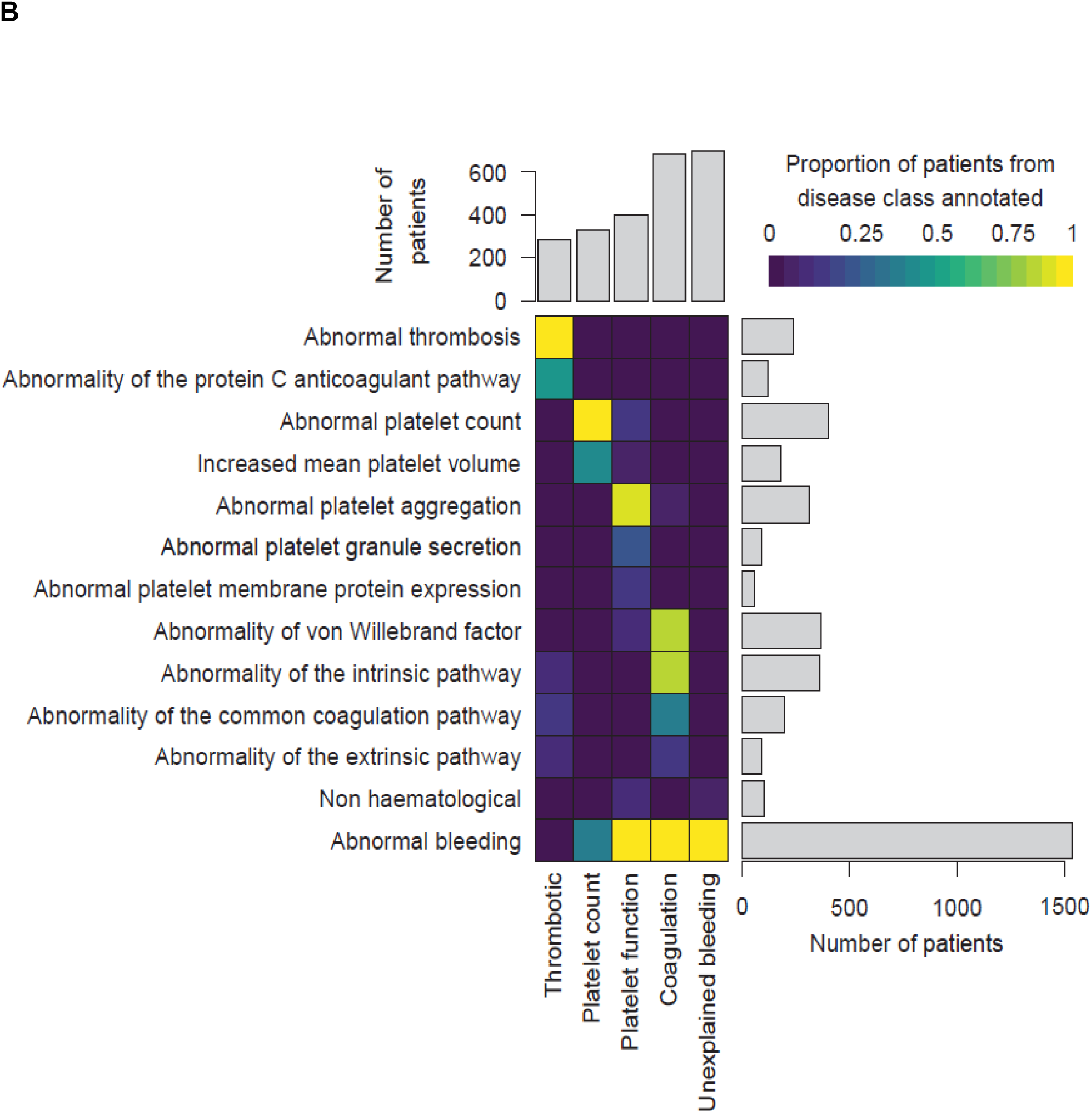
Classification of patients using clinical and laboratory phenotypes. (A) Classification of all patients from the ThromboGenomics, VIBB and PANE cohorts into one of five disease classes; thrombotic, platelet count, platelet function, coagulation and unexplained bleeding. (B) Representative HPO codes for patients characterised in each of the five disease classes.

Most of the patients categorised to the thrombotic class were referred because of reduced Protein C or Protein S levels as indicated by the HPO term ‘Abnormality of the protein C anticoagulation pathway’ (Figure 1B). Patients with platelet count disorders were generally referred due to (macro)thrombocytopenia. Platelet function abnormalities were diverse, including defects in aggregation, reduced platelet membrane protein expression (particularly GPIb/IX/V and GPIIb/IIIa) and reduced granule secretion. Coagulation defects included patients with reduced VWF levels or abnormal coagulation parameters of the intrinsic, extrinsic and common pathways. HPO terms coding abnormal phenotypes outside the blood system were appended to 41 (1.7%) of patients. These terms related to eyes (ocular albinism), hearing (deafness), skeletal (abnormal radius, joint disorders) and kidney (functional insufficiency). Across the three study groups 698 patients with bleeding symptoms and normal hemostasis test results were classed as unexplained bleeding (Figure 1B). Nearly half (49.1%) of the patients in this class were characterised by the presence of both spontaneous and trauma-related bleeding symptoms, a similar distribution as found for the coagulation and platelet function disease class (Supplemental Figure 2). Chronic mild thrombocytopenia is generally not accompanied by spontaneous bleeding.^18^ However, 67.7% of the patients in the platelet count class presented with spontaneous bleeding, strongly suggesting an enrichment of this type of ‘thrombocytopenia with bleeding’ referrals for the ThromboGenomics test.

### Performance of the ThromboGenomics HTS test

The previously reported validation of the ThromboGenomics HTS test (TG.V1) used 296 samples from patients, with and without previously known disease causing variants, sequenced for 63 TIER1 genes.^4^ Here we sequenced 1,330 and 1,060 index patient samples with the TG.V2 and TG.V3 tests targeting 80 and 96 TIER1 genes, with a region of interest (ROI) of 0.222 Mb and 0.275 Mb respectively (Supplemental Table 3). Since validation, the analysis method for calling SNVs and short (<50 base pairs [bp]) insertion/deletions (indels) has undergone minor modifications, however, the detection of CNVs has substantially been improved.^19^ In short, sequencing read depth is computed over 500 bp elements, to improve the sensitivity for the detection of shorter CNVs, and an optimised reference set of data obtained from 10 samples has been generated, using genetically unrelated individuals (Supplemental Information). Despite the increase of the ROI and increased multiplexing of samples, the read coverage has remained high, with 99.99% and 99.98% of the ROI with a read depth exceeding 30x for TG.V2 and TG.V3 tests, respectively. For each sample, an average of 146.7 and 190.9 SNVs, 9.4 and 11.2 indels and 0.20 and 0.21 CNVs were identified by the TG.V2 and TG.V3 tests, respectively. The proportions of variant types identified did not differ between test versions (Supplemental Figure 3). For all samples, on average 4.5 variants were prioritised for interpretation by the MDT (Supplemental Table 1).

### Diagnostic rates and validation of recently discovered TIER1 genes

Prioritised variants were reviewed by the MDT in the context of the disease incidence, variant frequency, assigned HPO terms and family history. Variants were reported with pathogenicity and the contribution to the patients phenotype (full or partial). Variants for recessive BPDs were reported if present in the homozygous or compound heterozygous states. VUS were reported if the MDT predicted a future Likely Pathogenic status with additional evidence from cosegregation and functional studies. Screening of 2,390 index patients resulted in an overall diagnostic rate of 37.5% by reporting a total of 1,039 variants in 897 index patients (Figure 2A). Most reported variants (88.5%) are rare (<0.01%) or absent in gnomAD (Supplemental Figure 4, Supplemental Table 6). There was a marked difference in diagnostic yield between the five classes; thrombotic, 49.1%; platelet count 48.0%; platelet function 27.0%; coagulation disorders 65.4% and unexplained bleeding 6.2% (Figure 2A). For thrombotic and coagulation disorders, 68.0% and 69.1% of variants reported were known variants that had previously been associated with disease, while for the platelet count and function disorders this proportion was lower; 43.5% and 26.4% respectively (Figure 2B). We reason that this difference reflects the fact that cataloging of pathogenic variants for coagulation disorders (especially VWD and Hemophilia A and B) began over three decades ago, whilst the majority of TIER1 genes for platelet disorders have only been identified over the past decade. Of the 329 patients within the platelet count class, 29 were referred under a working diagnosis of ‘immune thrombocytopenia refractory to treatment’. In 7 of these patients, variants were reported in genes known to be associated with thrombocytopenia (*ANKRD26, ETV6, ITGA2B, TUBB1*).

**Figure 2.**
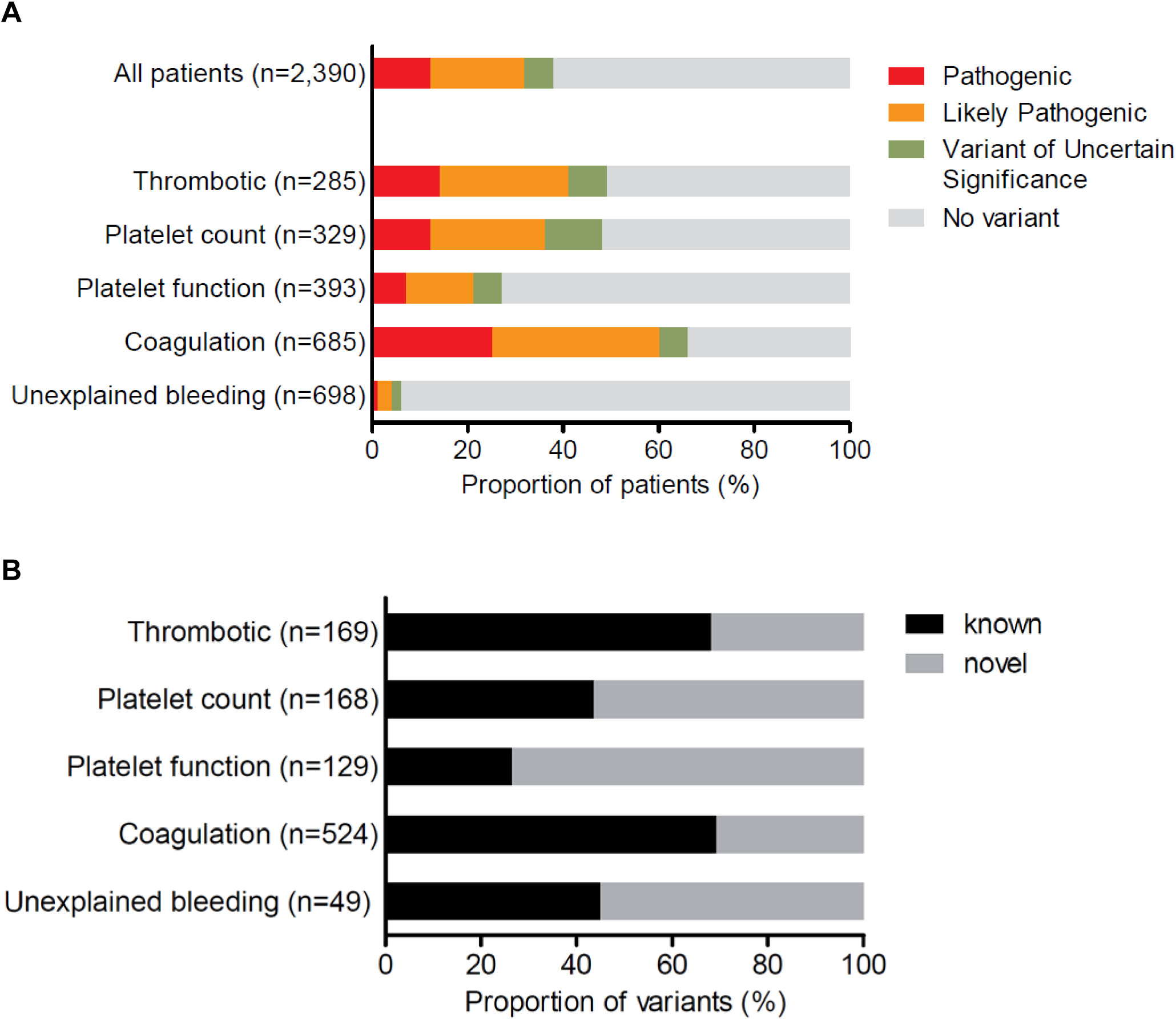
Diagnostic yield and proportion of novel variants by disease class. (A) Diagnostic yield of reported variants for 2,390 index patients for each of the five disease classes; thrombotic, platelet count, platelet function, coagulation and unexplained bleeding. For patients with more than one reported variant, the most pathogenic variant was used in this analysis (n = number of index patients). (B) Proportion of reported variants that were novel or known that were reported for patients in each disease class (n = number of variants).

The implementation of the criteria of the ACMG guidelines, instead of our ‘in-house’ criteria (Supplemental Table 4), impacted on variant interpretation for TG.V3. On comparing the collection of ThromboGenomics samples sequenced and analysed using either TG.V2 or TG.V3 (PANE and VIBB samples were sequenced using only TG.V2 and TG.V3 respectively), there was minimal difference in the total number of reported variants for each of the five classes of patients (Supplemental Figure 5). However, for all classes, an interpretation shift from Likely Pathogenic to VUS was noted. This change in variant interpretation was mainly explained by novel missense variants (Supplemental Figure 6). For patients tested using TG.V2, variants that were not known as disease associated were often deemed as Likely Pathogenic by the MDT. On introduction of the ACMG guidelines, these novel missense variants did not reach the threshold for designation as Likely Pathogenic and thus were labeled as VUS.

The genes with reported variants in index patients are ranked in Figure 3. For the thrombotic, platelet count and coagulation disease classes, over a quarter of patients had variants reported in just one gene; *PROS1, MYH9* and *VWF*, respectively. The fourth quarter of patients for each patient class had reports spread across at least 7 genes. All reported variants are summarised in Supplemental Table 6 and have been submitted to the ClinVar database.^20^

**Figure 3.**
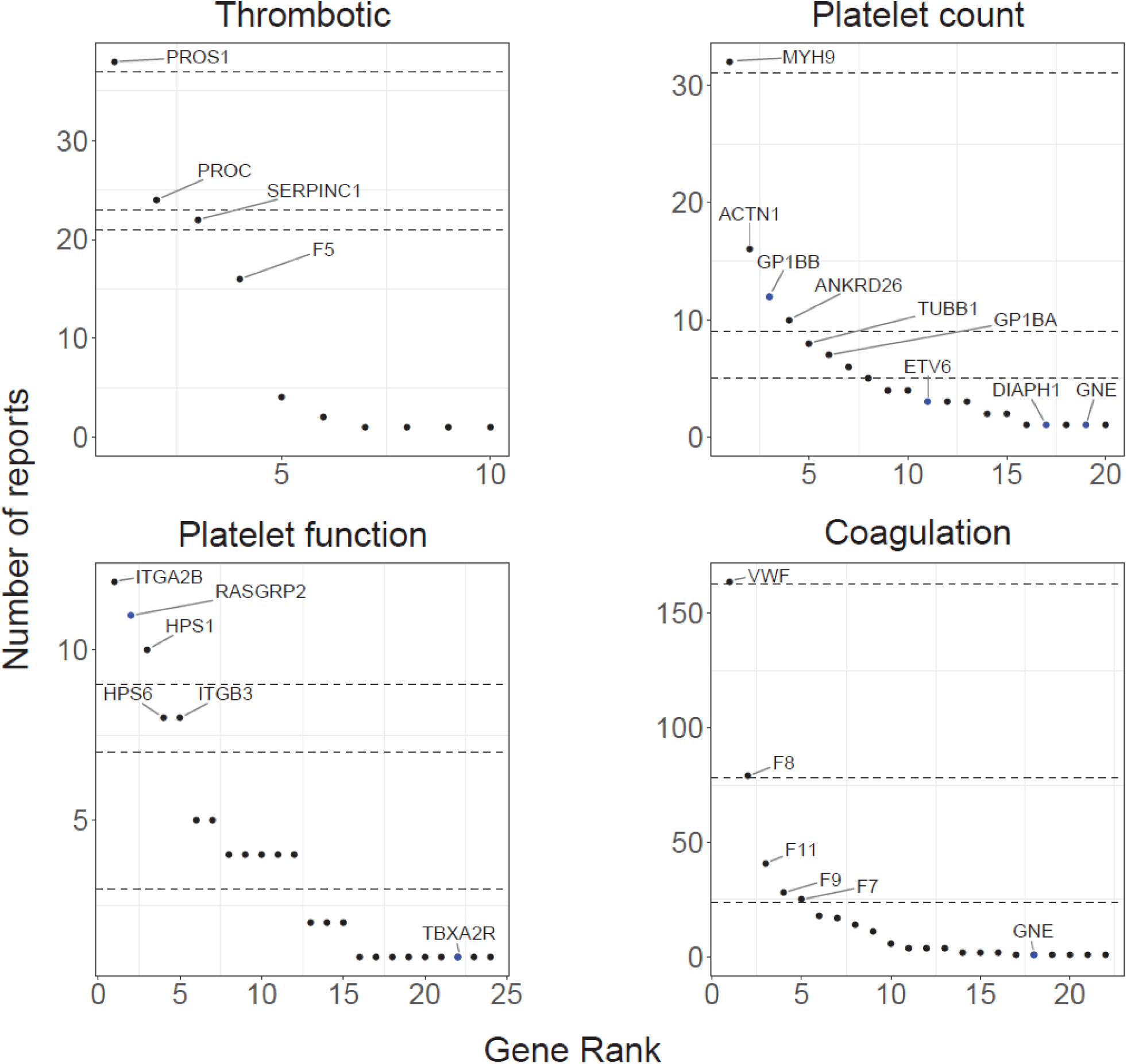
Gene ranking according to the number of reports per disease class. For each disease class, genes were ranked according to number of times they were reported. Dashed lines represents, from top to bottom, the 25, 50 and 75 quantiles. Recently discovered genes and changes of mode of inheritance are in blue.

Since TG.V1, 17 TIER1 genes associated with BPD have been included in TG.V2 and a further 16 more TIER1 genes in TG.V3. In addition to introducing known and recently discovered genes, also new modes of inheritance were added (detailed in Supplemental Table 3). Diagnostic reports for 41 patients have been issued with variants in one of the 19 recently discovered TIER1 genes or by applying a new mode of inheritance.

### Copy number variation, deep intronic variants and oligogenic findings

The analytical pipeline for the identification of CNVs has been modified with improved quality scoring and visualization tools. Overall, CNVs were reported in 40 patients, predicted to affect single exons (n=11), multiple exons (n=15) or whole genes (n=14) (Supplemental Table 6). These heterozygous deletions and duplications would not generally be detected using PCR-based and Sanger sequencing testing strategies. Raw sequencing reads of all potentially pathogenic CNVs were visually inspected by the MDT. In some instances, this revealed the presence of a complex CNV, including one inversion with breakpoints in introns 26 and 27 of *DIAPH1* resulting in an in-frame deletion of exon 26 and an inversion flanked by two deletions within *F8* causing severe hemophilia A (Supplemental Figure 7).^21^

Aside from the core dinucleotide splice sites at the 5’ and 3’ of introns, the lack of reliable prediction tools makes it difficult to determine the likely functional consequences of potential splicing altering variants. Outside the SnpEff annotated splice regions (8 bp), intronic variants were only prioritised if previously associated with disease. A deep intronic homozygous *ITGA2B* variant identified in one index patient with Glanzmann Thrombasthenia was validated using platelet RNA expression studies confirming alternative splicing and the absence of normal ITGA2B transcript (Supplemental Figure 8). These functional data, together with co-segregation analysis, in the pedigree of the index patient, resulted in a reclassification of this *ITGA2B* variant from VUS to Likely Pathogenic.

Of the 897 index patients where variants were reported, 773 had a single reported variant while 124 had at least two reported variants. The latter category were mainly patients with recessive diseases, however for 28 patients the variants were reported in two or more genes. For most of these examples, the variants identified were within first or second order interactors in the known canonical hemostasis pathways (Supplemental Table 7). For the thrombotic and coagulation classes, we identified 11 (3.9%) and 13 (1.6%) patients with oligogenic variants, respectively (Figure 4).

**Figure 4.**
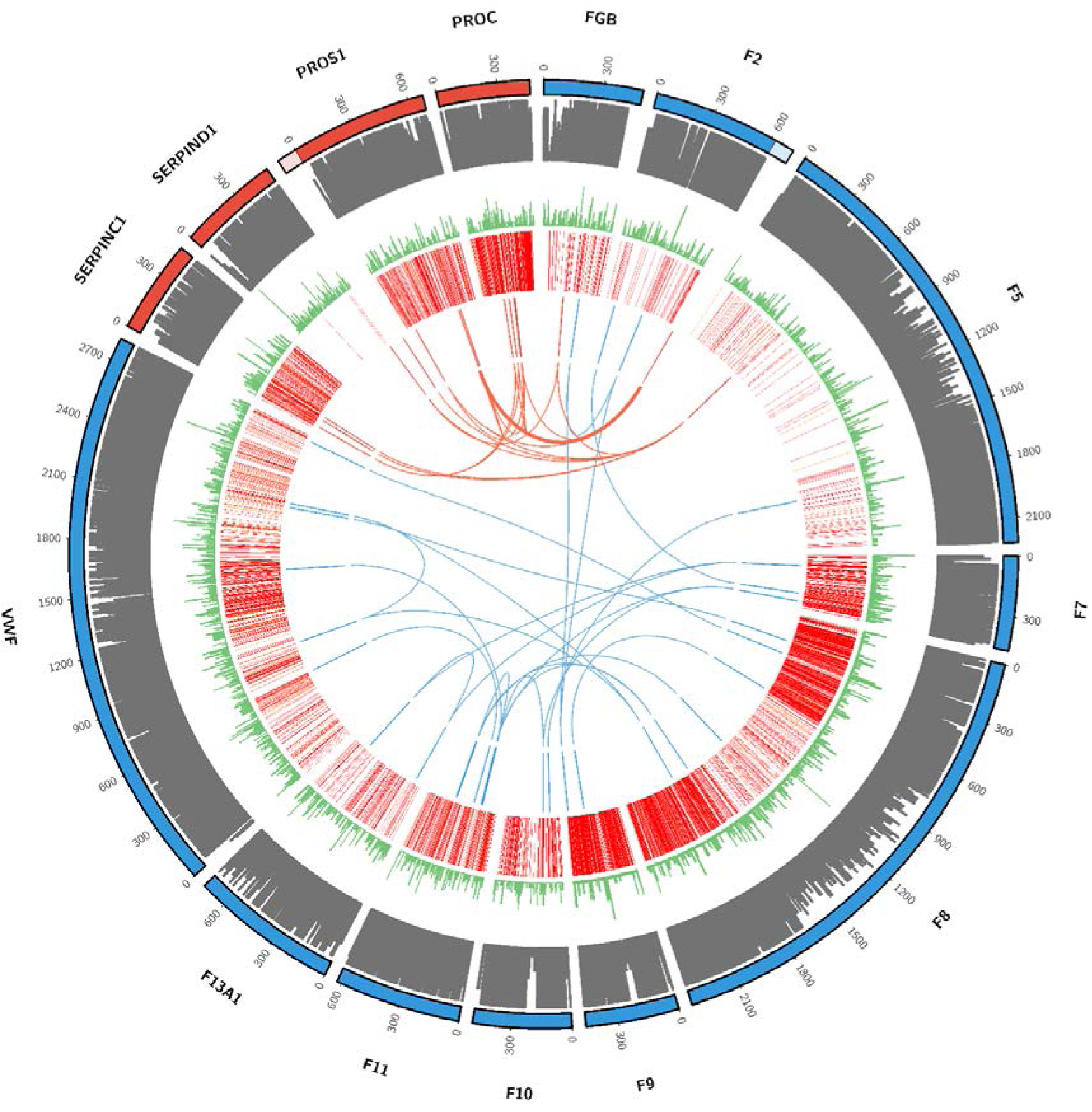
Oligogenic variants in patients with thrombotic (red) and coagulation (blue) disorders. From outside to in, track 1: Amino acid (AA) numbering with pro- and anti-coagulant proteins in blue and red, respectively; lighter shades denote sections of the F2 gene 3’ UTR or the PROS1 gene 5’ UTR; track 2: AA conservation scores calculated using ConSurf (http://consurf.tau.ac.il/2016/); track 3: variant frequency in gnomAD (minor allele frequency normalized scale to 1/106); track 4: Disease causing (DM, red) and potentially disease causing (DM?, orange) HGMD Pro variants; track 5 and arcs: reported variants in the 11 thrombotic (red) and 13 coagulation (blue) patients with oligogenic findings.

### Incidental findings

By sequencing a large number of patients for the TIER1 genes underlying known BPDs, we expected to observe incidental secondary findings. In four females, not referred for hemophilia carrier testing or known to have reduced factor levels, we identified carriership of Likely Pathogenic variants in *F8* or *F9*. In addition, in two patients we identified a heterozygous deletion of the RBM8A gene. A heterozygous *RBM8A* loss-of function variant (generally a deletion), if accompanied by a low frequency non-coding regulatory variant on the alternate allele, results in thrombocytopenia with absent radius (TAR) syndrome.^22^ These incidental secondary findings were reported to the referring clinician as they are actionable with respect to family planning. In contrast, sex chromosome aneuploidy, identified in three patients, was not reported in line with current guidance from the 100,000 Genomes Project.^23^

## Discussion

We evaluated the performance of a targeted HTS panel test for TIER1 genes in over 2,500 patients drawn from three distinct groups: (1) patients with a high likelihood of having an inherited BPD, (2) patients undergoing an extended preoperative assessment for bleeding risk and (3) patients with a bleeding disorder of unknown aetiology referred to a tertiary referral centre. After HPO coding of the clinical and laboratory phenotypes, patients were assigned to one of five diseases classes: thrombotic, platelet count, platelet function, coagulation or unexplained bleeding. DNA samples were sequenced with the ThromboGenomics HTS test and prioritised variants reviewed and classified by an MDT and reported to referring clinicians. The resulting data were used to assess the effectiveness of the HTS test, analytical pipeline and MDT variant interpretation in generating a conclusive molecular diagnosis. For patients of the thrombotic, coagulation and platelet count or function disease class, variants were reported for half (50.4%) of the 1,692 index patients. In contrast, variants were reported for 6.2% of the 698 index unexplained bleeding patients with normal hemostasis test results. Overall 20.1% (215) of variants reported were VUS, that require additional evidence including estimation of variant odds ratios using the results from large genotyped cohort studies, functional testing and cosegregation analysis. These data illustrate the excellent diagnostic yield of applying the ThromboGenomics test for patients with a high likelihood of having an inherited BPD.

Screening patients with the TG.V2 and TG.V3 HTS test presented several improvements compared to the TG.V1 test. Adding more TIER1 genes on TG.V2 and TG.V3 supported the molecular diagnosis of 41 patients and important genotype-phenotype association were observed for recently discovered BPD genes such as *DIAPH1^21^, ETV6^24^, GFI1B^25^*, GNE^26,27^ and *RASGRP2^28^* or for the alternative mode of inheritance recently reported for *GP1BB.^29^* In comparison to TG.V1, the detection of CNVs was optimised, resulting in the detection of CNVs in 40 patients, including previously unobserved deletions in 11 genes, indicating CNVs being an important variant class for all categories of BPDs. In addition, 2 novel duplications, 1 novel inversion and a complex CNV were also reported. Nevertheless, the optimised ExomeDepth method is sensitive to variable read depths, has a minimum resolution and cannot detect inversions or predict the location of duplicated regions. The introduction of a split-read or read-pair CNV calling method, alongside the ExomeDepth analysis method, may further improve CNV detection in the future.

Previously, Sanger sequencing of most BPD genes was performed with primer sets flanking intron/exon boundaries and therefore, the frequency of non-coding or silent variants, aside from those disrupting the immediate splice site (<+/− 8 bp from the exon boundary), that are associated with BPDs is unknown. With the use of HTS, deep intronic variants have been identified and apparent silent variants in BPD genes have been shown to alter splicing.^30–33^ Variants disrupting transcription regulatory motifs located in gene promoters and enhancer regions have also been associated with BPDs and with the recent mapping of endothelial and blood cell specific enhancers, it is likely that more variants located in these regions will be identified as associated with disease in the near future.^34–36^ Nevertheless, due to the challenges in interpretation of non-coding variants we have only targeted and prioritised intronic and regulatory variants if previously associated with disease. Therefore, a research analysis of novel deep intronic, silent and likely regulatory variants detected is required and such studies are best performed using whole genome sequencing data from well characterised patient cohorts.

A diagnostic HTS platform requires careful selection of diagnostic-grade TIER1 genes. The decision to include a gene to the BPD TIER1 list is made by the GinTH SSC of the ISTH. The designation of genes for diagnostic reporting involved the review of associated literature to evaluate if a gene is associated with disease in more than three independent pedigrees with convincing cosegregation data or in less than three pedigrees, but with strong functional evidence (mouse models or cell/protein studies) in addition to cosegregation data. In the future, this task will also be coordinated by the NIH supported Hemostasis/Thrombosis Clinical Domain Working Group in close partnership with the ISTH and ASH expert working groups.^37^

Our results indicate that when screening large number of patients with a HTS test that approximately 41% of reported variants are novel (Figure 2B), and following current guidelines, the majority of novel missense variants are reported as VUS (Supplemental Figure 6). International initiatives for sharing of sequence data generated for BPD patients and the sequencing of the genomes or exomes of large prospective population cohorts, like the UK Biobank, the Million Veteran Program and the 100,000 Genomes Project will lead to statistically robust approaches for the functional labelling of DNA variants, including the VUS reported in this study.^38–39^

We obtained evidence of how having reached a molecular diagnosis has influenced clinical management and counselling of patients and their close relatives. In 30 patients, variants were reported in the ANKRD26, ETV6 and RUNX1 genes leading to counseling and follow up due to the increased risk of hematological malignancies. In 7 patients with treatment refractory immune thrombocytopenia evidence was obtained of rare germline variants likely causing their condition. In six of these cases the VUS finding has prompted follow-up studies in the probands and their relatives to obtain additional evidence for pathogenicity. Finally, in 24 patients with thrombotic or coagulation disorders, we reported two or more variants in relevant TIER1 genes. We postulate that defects in hemostasis are due to the disruption of two interacting proteins in the known canonical pathways. There have been case reports of possible oligogenic architecture for thrombotic and coagulation disorders that add to the findings of this large single study. Together these justify further functional and genetic follow-up studies to provide patients with a molecular diagnosis and better estimates of risk to offspring.

All together we report on the results of the largest gene panel sequencing study of patients with possible inherited BPDs. We included 698 individuals with unexplained bleeding disorders with normal hemostasis test results and as per our hypothesis, we observed a low diagnostic rate for this group. It is possible that the propensity of bleeding in these patients is the result of the aggregation of a large number of small effects emanating from common variants at hundreds of loci which are modifying the overall effectiveness of the hemostasis system. Estimation of polygenic risk scores has recently been reported for several common diseases such as coronary artery disease, type 2 diabetes and breast cancer^40–43^.

In conclusion the ThromboGenomics test is a valuable addition to the diagnostic algorithm for patients with a high likelihood of having an inherited BPD. The results provide clinicians with a molecular diagnosis for approximately half of patients allowing for more precise prognostication and management of disease and, with cascade testing, better informed counselling of patients and their close relatives.

## Data availability

Variants reported as pathogenic, likely pathogenic and variants of uncertain significance have been deposited into ClinVar under accession number xxxx.

## Acknowledgments

This study makes use of data generated by the NIHR BioResource. A full list of investigators who contributed to the generation of the data is available from https://bioresource.nihr.ac.uk/researchers/researchers/acknowledgement/. We gratefully acknowledge the participation of all NIHR BioResource volunteers, and thank the NIHR BioResource centre and staff for their contribution. Funding for the project was provided by the National Institute for Health Research (NIHR) under grant number RG65966. The authors would like to thank the Locus Reference Genomic team (LRG) for their assistance in curating gene transcripts. Research in the Ouwehand laboratory receives funding from the British Heart Foundation, European Commission (TrainMALTA), International Society on Thrombosis and Haemostasis, Medical Research Council, NHS Blood and Transplant and the Rosetrees Trust. The Vienna Bleeding Biobank was supported by an unrestricted grant of CSL Behring. K.D. is supported as a NHS HSST trainee by Health Education England. K.F. is supported by the Research Council of the University of Leuven (BOF KU Leuven, Belgium; OT/14/098) and by an unrestricted grant of SOBI.

## Authorship

Contributions: D.D., J.S. and C.T. performed experiments; K.M., M.V., O.S., S.V.V.D., R.M., S.T., L.D., N.G., D.G., M.H., A.T., C.J.P., E.T. analysed data; J.G., S.H., M.W.B., N.C., S.P., S.R-V., E.S., S.S., E.S., W.T., I.S., Y.M.C.H. and I.P., collected samples and provided clinical support; N.A.H., H.M., and S.A. provided clinical support; K.D., W.H.O., M.A.L., A.D.M., K.G. and K.F. are members of the ThromboGenomics MDT and designed the study and wrote the manuscript, which was reviewed by all remaining authors.

## Supplemental Information

Diagnostic high-throughput sequencing of 2,390 patients with bleeding, thrombotic and platelet disorders

### Patient cohorts

#### ThromboGenomics cohort

International recruitment of 1602 index patients with mostly known or suspected inherited bleeding, thrombotic or platelet disorders following previously described criteria used for the validation of the ThromboGenomics HTS test (Supplemental Table 1). ^1^ Patients were recruited or referred to this study through two routes.

Patients recruited to the bleeding and platelet disorder arm of the NIHR-BioResource Rare Diseases study with a suspected disease etiology were sequenced using the HTS test as a pre screen prior to whole genome sequencing. Patients provided written consent according to the study ethical approval (East of England Cambridge South national research ethics committee (REC) reference 13/EE/0325). Obtaining consent from overseas patients was the responsibility of the respective principal investigators at the enrolling hospitals. Material and transfer agreements were applied to regulate the exchange of samples and data between the donor institutions and the University of Cambridge.

Patients were referred for clinical testing and consented by the referring clinician, according to local clinical practice. For UK patients, a UKHCDO approved patient information leaflet and consent form was provided for use. Consent for genetic testing was also obtained from patients and a small number of relatives and carriers (Supplemental Table 1).

Patients referred for clinical testing with severe hemophilia A were tested for the pathogenic intron 1 and intron 22 inversions in the F8 gene using inverse shifting-polymerase chain reaction prior to testing using the ThromboGenomics test.^2^

All likely pathogenic and pathogenic variants in patients referred for clinical testing have been confirmed using Sanger sequencing or Multiplex ligation-dependent probe amplification (MLPA) copy number variation (CNV) analysis in the Cambridge University Hospitals Genetic Laboratories.

### Preoperative screening for mild bleeding disorders cohort (PANE)

Individuals were identified using responses to a pre operative anesthesiology bleeding questionnaire when admitted for elective surgery at Maastricht University Medical Centre (MUMC). Patients were invited to join the PANE study if they reported one or more bleeding symptoms and were over the age of 18, had no anaemia and were not using drugs that could interfere with hemostasis.^3,4^ Ethical approval was obtained from the Medical Ethics Committee of MUMC (NL38767.068.11/METC11–2–096). Written informed consent was obtained from all patients. During the study visit, bleeding symptoms of the participants were evaluated by an experienced hematologist. Blood was drawn and subjected to a panel of 30 laboratory tests (Supplemental Table 2). Using local reference values, HPO codes were appended to patients with abnormal test parameters and clinical bleeding symptoms. HTS was performed for 212 unrelated study subjects that included 193 index patients and 19 subjects that did not have any bleeding symptoms or clinical test results that indicated a BPD-related HPO code (Supplemental Table 1).

### Vienna Bleeding Biobank cohort (VIBB)

The VIBB study was established in collaboration with the MedUni Wien Biobank (Department of Laboratory Medicine, Medical University of Vienna, Austria, www.biobank.at) as a single-centre study.^5^ Patients who were referred to the hemostasis outpatient department for investigation of a mild to moderate bleeding disorder were recruited into the The Vienna Bleeding Biobank (VIBB) study. The Ethics Committee of the Medical University of Vienna approved the project (EK No 603/2009 and 039/2006). All patients gave written informed consent before inclusion in the VIBB study. On recruitment to the study, blood was drawn and subjected to a panel of 42 laboratory tests (Supplemental Table 2). Using local reference values, HPO codes were appended to patients with abnormal test parameters and clinical bleeding symptoms. HTS was performed for 594 index patients (Supplemental Table 1).

### ThromboGenomics HTS BPD test design

ROCHE NimbleGen SeqCap capture baits (ROCHE NimbleGen, Inc. Madison, WI USA) were designed to target all consensus coding sequences (CCDS) of the ThromboGenomics genes, the first and last 100 bp of introns, 5’ and 3’ UTRs, regions 1,000bp upstream of the transcription start site and the position of known pathogenic variants (see below). Regions of four allosome genes (SRY, TSPY1, AMELY, AMELX) were included to assign genomics sex. For TG.V3, capture baits were also included for a panel of 10,000 common SNVs to calculate ethnicity and relatedness estimates.

### Known variants

A curated list of known disease associated variants were included in the HTS test design and for variant prioritisation.

At the time of design, all known Human Gene Mutation Database (HGMD) variants associated with the BPD genes were included in the design.^6^ For TG.V2, 9,280 variants were included (HGMDPro 2015.2) and for TG.V3, 11,442 variants were included (HGMDPro 2016.4).

A set of known variants was used to prioritise the variants identified in patient samples (see below). All variants in HGMD_PRO_2017.2 were used to perform the prioritisation analysis of all TG.V2 samples. HGMD_PRO_2017.4 variants were used for the prioritisation of TG.V3 samples alongside a curated list of variants from disease specific databases. These included; 2,498 variants, with a gnomAD minor allele frequency (MAF) <0.25, from the F7, F8, F9 and VWF gene EAHAD databases (http://www.eahad-db.org/, accessed February 2017); 438 MYH9, GP1BA, GP1BB, GP9, and WAS variants with a MAF <0.01 from the LOVD databases^7^ (accessed May 2017); 203 ITGA2B and ITGB3 variants from the Glanzmann Thrombasthenia Database (https://glanzmann.mcw.edu/, accessed May 2017). In addition, all variants previously reported by ThromboGenomics were used for prioritisation.

### Library preparation, enrichment and sequencing

DNA samples were processed as previously described with minor modifications.^1^ In short, samples were processed in batches with 500ng of each sample fragmented using a Covaris E220 (Covaris Inc., Woburn, MA, USA). Samples were processed using the ROCHE KAPA HTP Library Preparation kit (Roche Diagnostics Ltd., Burgess Hill, UK). DNA libraries were captured using ROCHE NimbleGen SeqCap ThromboGenomics capture baits (ROCHE NimbleGen, Inc. Madison, WI USA). Final libraries were quantified, samples pooled and sequenced using an Illumina Hiseq 4000 sequencer, 150 base pair (bp) paired-end (PE) run.

Initially for TG.V2, 48 samples were multiplexed per sequencing reaction. In an effort to increase sample throughput and reduce costs, methods were adjusted to multiplex 96 samples. At the same time, modifications were made to increase the DNA library fragment size with the aim of increasing read coverage in poorly performing regions. DNA was fragmented to obtain an average insert size of 220 bp for TG.V2 and 350 bp for TG.V3.

### Variant calling

Single nucleotide variants (SNVs) and short insertions or deletions (INDELs) were called using GATK 3.3 using GRCh37.^8^ HaplotypeCaller in a single sample mode and filtered using the following VariantFiltration expressions “MQ< 40.0 || QD < 2.0 || FS > 100.0” for SNVs and “FS > 200.0 || QD < 2.0 || ReadPosRankSum < −20.0” for INDELs. Variants were merged into multi-sample VCF files. SNVs and INDELs were annotated with their predicted impact against Ensembl 75, presence in the human gene using SnpEff 4.0.^9^

### Relatedness and ancestry estimation

A panel of 10,000 SNVs were incorporated into the design of TG.V3 to estimate the degree of relatedness between individuals and to categorise an individual’s ancestry into European, African, East Asian, South Asian or Other. These variants were selected from a larger panel of SNVs recommended by ROCHE for this purpose (personal communication, Todd Richmond, Roche Sequencing Solutions). Principal component analysis of samples from the 1000 Genomes Project with known ethnicity were used to generate a reference data set. Patient samples were then compared to this reference to estimate ancestry.

Relatedness was estimated using the 10,000 common SNVs using the PC-Relate function from GENESIS R package. For each sample pair, a relatedness score was calculated, ranging from 0 to 1.

### Copy Number Variation

CVNs were called using a custom pipeline based on the ExomeDepth R-package (version 1.1.10).^10^ ExomeDepth makes a copy number gain or loss call in a specified genomic interval by comparing the read depths in a sample and an optimised reference set of other samples. Our customisation reduces false negative and positive calls by specifically defining a set of ten unrelated reference samples. To detect small CNVs within large exons, genomic intervals of no more than 500bp were used to calculate read depth. Modifications were also made to the ExomeDepth read counting method to avoid inflation caused by reads overlapping two adjacent genomic intervals. CNVs observed in more than 10% of samples within a batch were filtered out as technical artefacts or common CNVs.

### Region of interest

The region of interest (ROI) for variant prioritisation was defined as:

– all coding regions for each curated gene transcript
– +/− 15 bps into the introns
– 5’ and 3’ UTRs sequences
– the position of all known variants at the time of the panel design

### Variant prioritisation

For each patient sample, variants identified were prioritised to provide a list of potentially pathogenic variants for interpretation by the multi-disciplinary team (MDT).

Variants were prioritised if:

– predicted to have a moderate or a high impact effect according to SnpEff
– within the snRNA gene RNU4ATAC
– located at the same nucleotide position as a known variant with a gnomAD MAF <0.025 or
– novel with a gnomAD MAF <0.001

Variants were not prioritised if they had >3 alternate alleles (to guard against sequencing errors in repetitive regions) or if observed in the HTS samples with a frequency >=10% (to remove systematic artifacts).

### Problematic regions

A number of regions are not well covered by aligned sequence generated using the HTS test and bioinformatic pipeline. A region in the UTRs of the ORAI1 gene has poor sequencing coverage, although only 6bp have a read depth less than 20x (GRCh37:Chr12:122064774–122064779).

Exon 26 of the VWF gene has poor aligned sequence coverage due to a homologous region within VWFP1, an unprocessed pseudogene on chromosome 22 (<20x aligned reads at GRCh37:Chr12:6131938–6132010). Raw sequencing reads that align to this region are manually inspected in patients with an indication of von Willebrand Disease to identify any possible variants within this exon. Any potential pathogenic variants are confirmed using PCR primers specific to the VWF gene sequence before issuing a report.

## Supplemental Tables and Figures

**Supplemental Table 3** - excel sheet

Genes and disorders included in the ThromboGenomics HTS test. TIER1 genes sequenced using the ThromboGenomics HTS panel including disease category, HGNC-approved gene symbol, gene name, disorder tested, mode of inheritance, reporting transcript, LRG accession and version of test. AD, autosomal dominant; AR, autosomal recessive; XLR, X-linked recessive. *Gene without sufficient evidence for TIER1 status as of July 2018.

**Supplemental Table 6** - excel sheet

Variants reported in index patients using the ThromboGenomics HTS test.

Single nucleotide variants and small insertion-deletions are in a separate tab to the CNVs.

**Supplemental Table 1.** Summary of the subjects from the ThromboGenomics, PANE and VIBB collections sequenced using the HTS test

|  | Cohort size |  |  |  |  | 2390 Index patients<br>(prioritised variants) |  | Diagnostic<br>yield |
| --- | --- | --- | --- | --- | --- | --- | --- | --- |
| | Index<br>cases | Hemophilia<br>carrier query<sup>\$</sup> | Affected<br>relatives | Non affected<br>relatives<sup>&</sup> | Non affected<br>individuals<sup>#</sup> | TG.V2 | TG.V3 | % patients |
| **TG** | 1,602 | 18 | 98* | 33* | - | 1136 patients<br>(4.75 variants) | 466 patients<br>(4.70 variants) | 52.2% |
| **PANE** | 193 | - | - | - | 19 | 193 patients<br>(3.28 variants) | - | 4.6% |
| **VIBB** | 594 | - | 7<br>(mother<br>child pairs) | - | 4 | - | 594 patients<br>(4.21 variants) | 10.4% |

**Supplemental Table 2.**
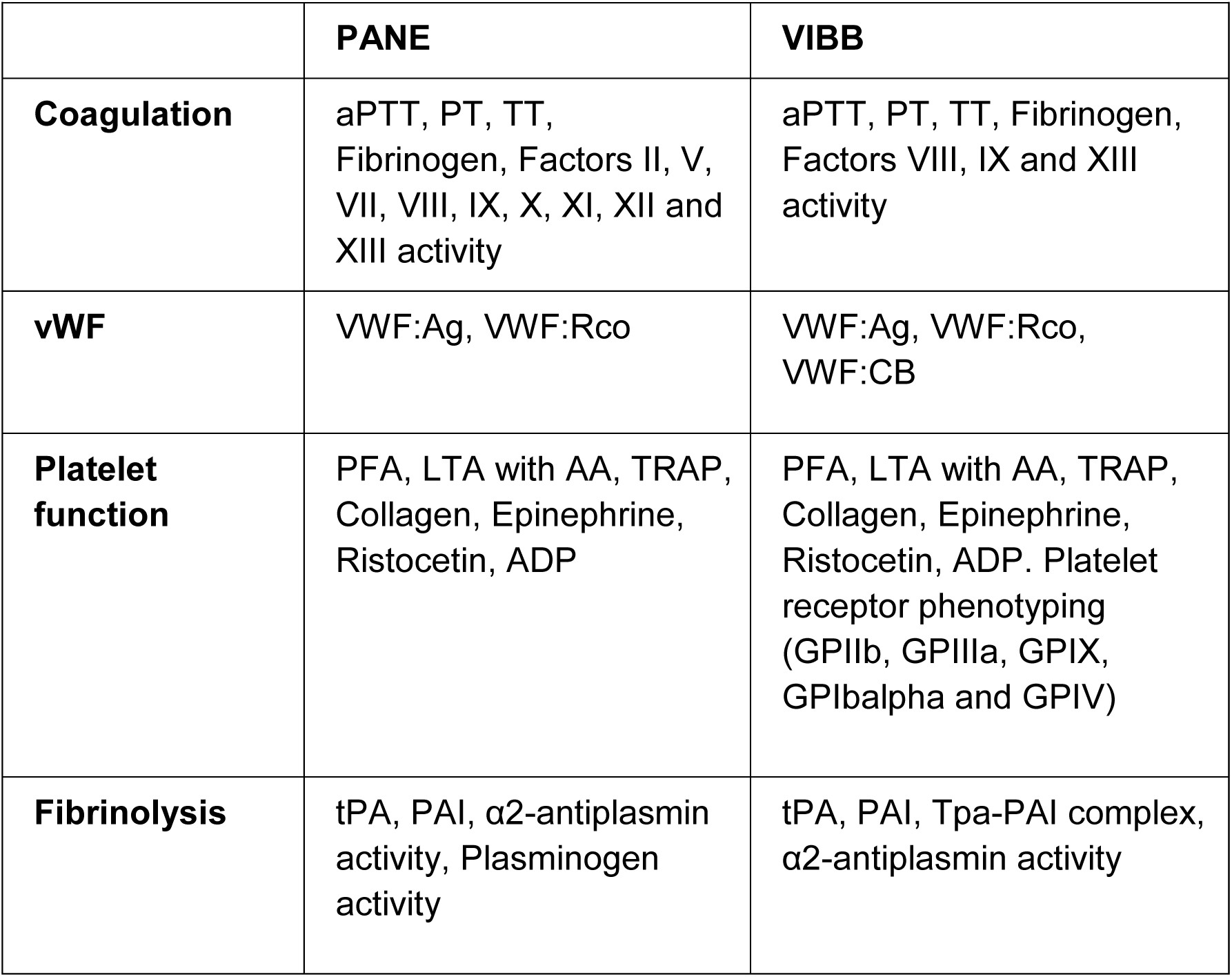
Key laboratory tests performed in the PANE and VIBB study subjects.

**Supplemental Table 4.** Variant interpretation guidelines used by ThromboGenomics multi-disciplinary team for study participants sequenced using TG.V2.

| Variant classification | Criteria |
| --- | --- |
| Pathogenic | Variant that has been described in at least 3 unrelated index patients that have been reported in:<br>- HGMD as disease mutation (DM)<br>- F7/F8/F9/VWF EAHAD variant database (http://eahad-db.org)<br>- ThromboGenomics BPD variant dataset |
| Likely Pathogenic | Variant that has been described in <3 unrelated index patients but has been reported in:<br>- HGMD as DM<br>- F7/F8/F9/VWF EAHAD variant database<br>- ThromboGenomics BPD variant dataset<br>OR<br>Variant that is absent from control datasets (1000G, ExAC and UK10K sequencing data) and present in a gene that strongly matches the clinical and laboratory phenotype for the patient.<br>OR<br>Variant that is absent from control datasets (1000G, ExAC and UK10K sequencing data) and predicted to cause a loss of function |
| Variant of uncertain clinical significance (VUS) | Variant that has a low minor allele frequency (MAF) or is absent from control datasets (1000G, gnomAD and UK10K sequencing data) and present in a gene that matches the clinical and laboratory phenotype for the patient |

**Supplemental Table 5.** Summary of differences between the ThromboGenomics TG.V2 and TG.V3 HTS tests, analysis methods and variant interpretation.

|  | **TG.V2** | **TG.V3** |
| --- | --- | --- |
| **Panel content** | 19 Coagulation genes<br>8 Thrombotic genes<br>53 Platelet genes<br>HGMDPro 2015.2 variants | 21 Coagulation genes<br>9 Thrombotic genes<br>66 Platelet genes<br>HGMDPro 2016.4 variants<br>10,000 SNVs |
| **Region of Interest (ROI)** | 0.222 Mb | 0.275 Mb |
| **Methods** | DNA fragmentation – 220 bp<br>Multiplex – 48 samples | DNA fragmentation – 350 bp<br>Multiplex – 96 samples |
| **Variants used in prioritisation analysis** | HGMD_PRO_2017.2 | HGMD_PRO_2017.4<br>Curated known variants |
| **Variant interpretation** | In house ThromboGenomics<br>MDT criteria (see<br>Supplemental Table 4) | ACMG guidelines* |

\* Richards S, Aziz N, Bale S, et al. Standards and guidelines for the interpretation of sequence variants: a joint consensus recommendation of the American College of Medical Genetics and Genomics and the Association for Molecular Pathology. *Genet Med.* 2015;17(5):405-424.

**Supplemental Table 7.** Details of index patients with oligogenic findings. VUS – Variants of Uncertain Significance.

| Disease class | Index patient | Gene 1 | Gene 2 | Gene 3 |
| --- | --- | --- | --- | --- |
|  |  | Variant (heterozygous/homozygous) – Variant classification |  |  |
| Thrombotic | 1 | F5 (het)- Pathogenic | PROS1 (het)- Likely Pathogenic |  |
|  | 2 | F5 (het)- Pathogenic | PROS1 (het)- VUS |  |
|  | 3 | SERPIND1 (het)- VUS | FGB (het)- VUS | F5 (het)- Pathogenic |
|  | 4 | SERPINC1 (het)- Likely Pathogenic | F5 (het)- Pathogenic |  |
|  | 5 | F2 (het)- Likely Pathogenic | PROS1 (het)- Likely Pathogenic |  |
|  | 6 | SERPINC1 (het)- Pathogenic | F5 (het)- Pathogenic |  |
|  | 7 | PROS1 (het)- Likely Pathogenic | PROC (het)- Likely Pathogenic | F2 (het)- Pathogenic |
|  | 8 | F2 (het)- Pathogenic | PROS1 (het:variant 1)- Likely Pathogenic<br>PROS1 (het:variant 2)- Likely Pathogenic |  |
|  | 9 | F2 (het)- Pathogenic | PROS1 (het)- Pathogenic |  |
|  | 10 | PROC (het)- Likely Pathogenic | PROS1 (het)- Pathogenic |  |
|  | 11 | PROC (het)- Likely Pathogenic<br>PROC (het)- Likely Pathogenic | SERPINC1 (het)- VUS |  |
| Coagulation | 1 | F8 (homo) - Pathogenic | F9 (homo)- VUS |  |
|  | 2 | F11 (het:variant 1)- Likely Pathogenic<br>F11 (het:variant 2) - Likely Pathogenic | F8 (het)- VUS |  |
| **Coagulation** | **3** | F2 (het)- Likely Pathogenic | FGB (het)- Likely Pathogenic | F9 (het)- Pathogenic |
|  | **4** | VWF (het)- Likely Pathogenic | F11 (het)- Likely Pathogenic |  |
|  | **5** | F2 (het)- Likely Pathogenic | F7 (het)- Likely Pathogenic |  |
|  | **6** | VWF (het:variant1)– VUS<br>VWF (het:variant2)– VUS | F8 (het)- VUS |  |
|  | **7** | VWF (het)- VUS | F8 (het)- Pathogenic |  |
|  | **8** | F11 (het)- VLP Likely Pathogenic | F10 (het)- VUS | F7 (het)- VUS |
|  | **9** | F11 (het)- Pathogenic | VWF (het)- Likely Pathogenic |  |
|  | **10** | F8 (homo)- Likely Pathogenic | VWF (het)- VUS |  |
|  | **11** | F5 (het)- Likely Pathogenic | F10 (het)- VUS |  |
|  | **12** | F8 (homo)- VUS | F11 (het)- Likely Pathogenic |  |
|  | **13** | F13A1 (het:variant 1) - Likely Pathogenic<br>F13A1 (het:variant 2) - Likely Pathogenic | F7 (het)- VUS |  |
| **Coagulation and Platelet function** | **1** | VWF (het)- Likely Pathogenic | P2RY12 (het)- Likely Pathogenic |  |
|  | **2** | VWF (het)- Likely Pathogenic | GP1BB (het)- VUS |  |
| **Coagulation and Platelet count** | **1** | VWF (het)- Pathogenic | GATA1 (het)- VUS |  |
|  | **2** | F11 (het)- VUS | TUBB1 (het)- VUS | MYH9 (het)- VUS |
| **Bleeding** | **1** | SERPINF2 (het)- VUS | THBD (het)- VUS |  |

## Supplemental Figures

**Supplemental Figure 1.**
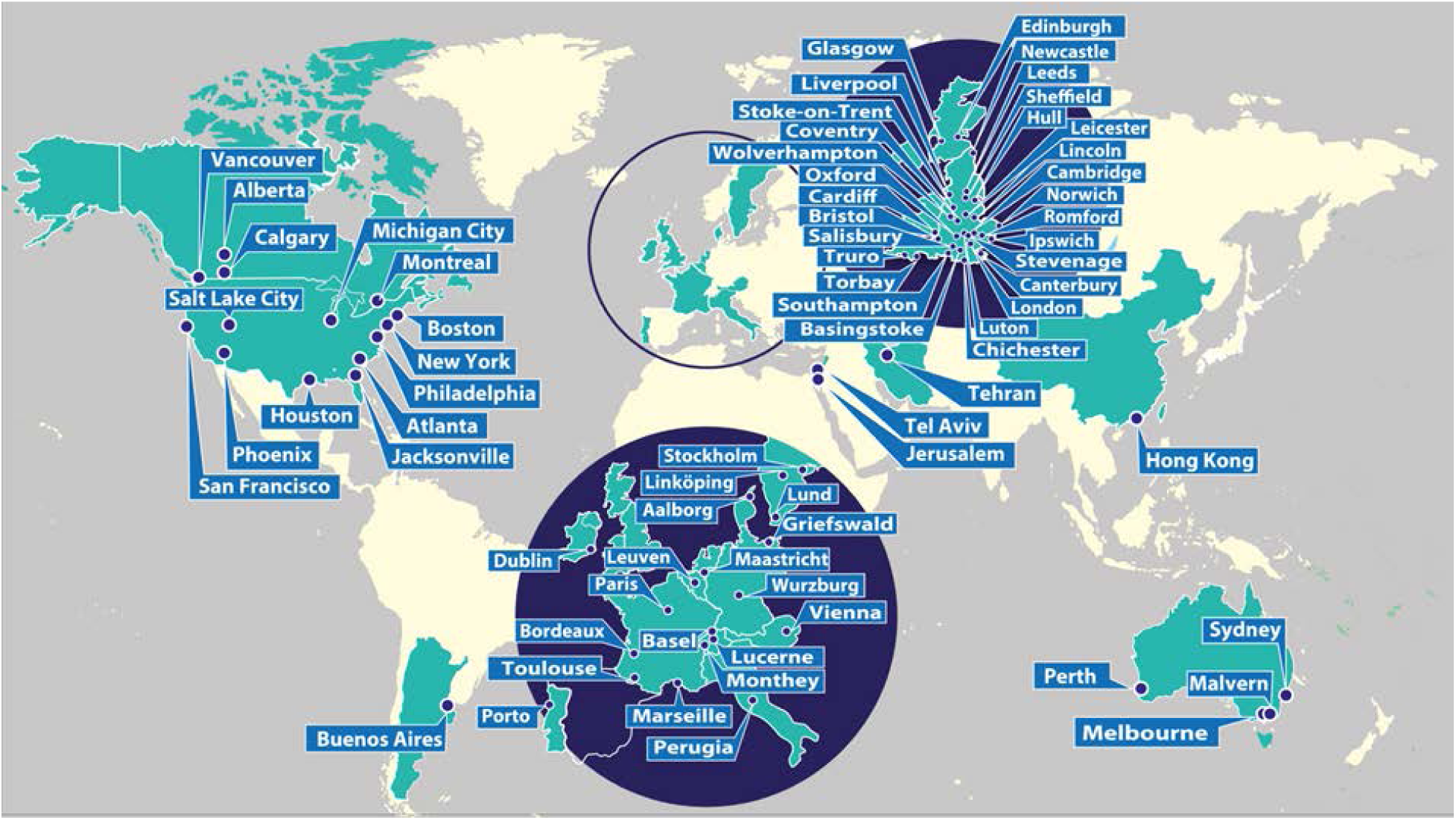
A global map with the location of 72 UK and 46 non-UK hospitals which referred samples to ThromboGenomics.

**Supplemental Figure 2.**
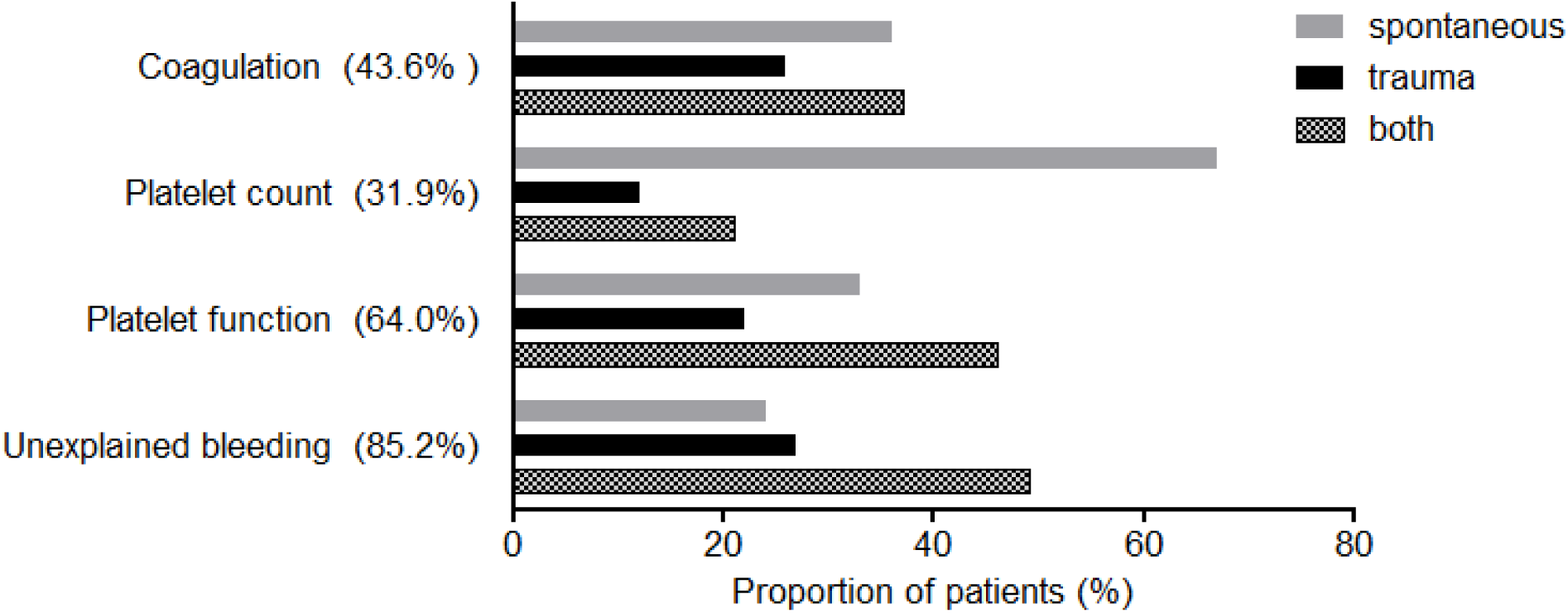
Distribution of bleeding symptoms present in the different disease classes. The figure represents information from all patients with assigned HPO codes for ‘clinically significant bleeding events’ that were present in at least 10 patients. The proportion of patients per type of bleeding is on the horizontal axis. The HPO term for menorrhagia was excluded when selecting patients for this analysis. Between brackets are the proportion of patients within each disease class with bleeding phenotypes used in this analysis.

**Supplemental Figure 3.**
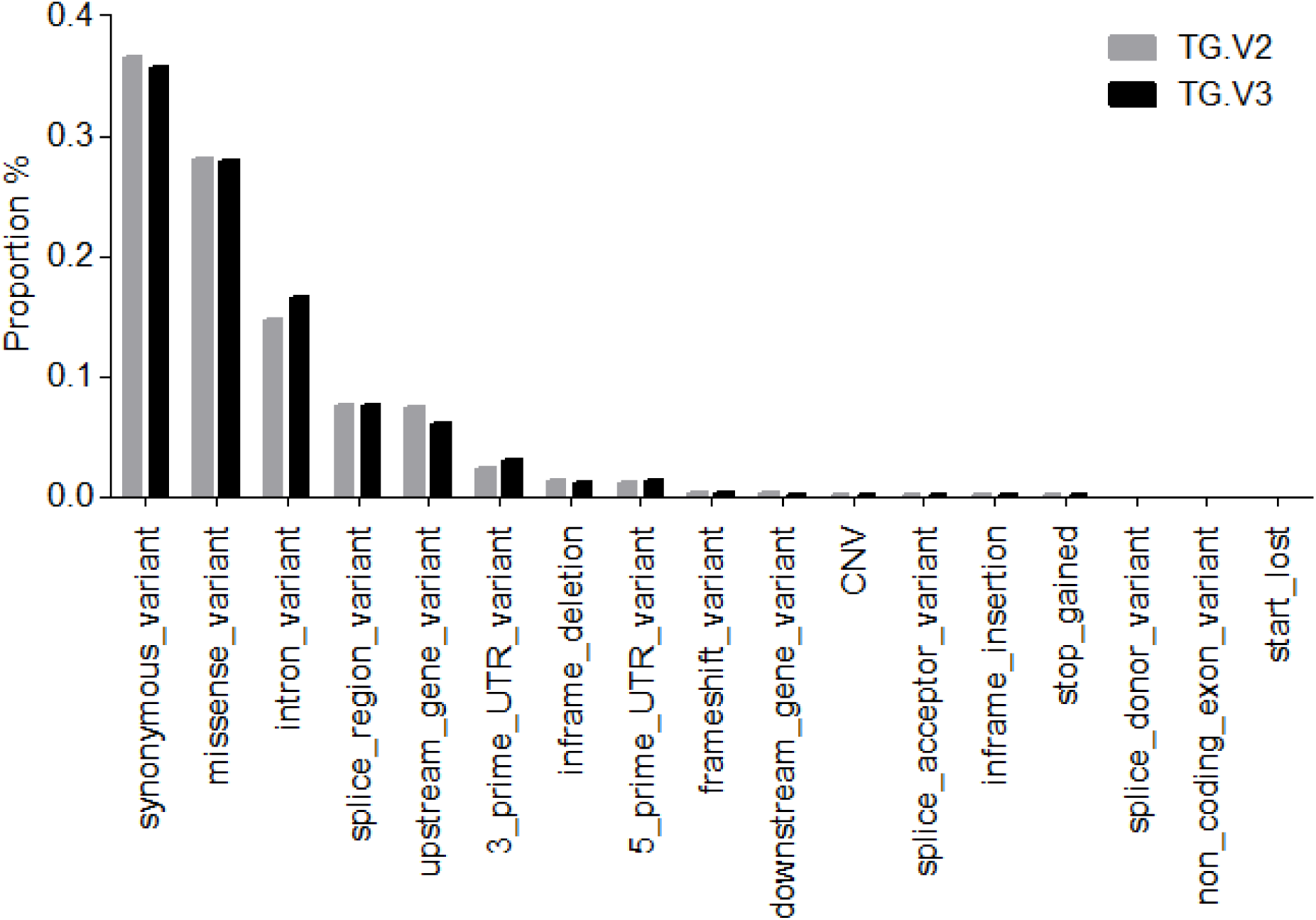
The average proportion of variants and predicted effects identified in patients sequenced using TG.V2 or TG.V3. Only variants within the region of interest (ROI) were included. The average number of variants within the ROI for each patient was 156.3 and 202.4 for TG.V2 and TG.V3 respectively.

**Supplemental Figure 4.**
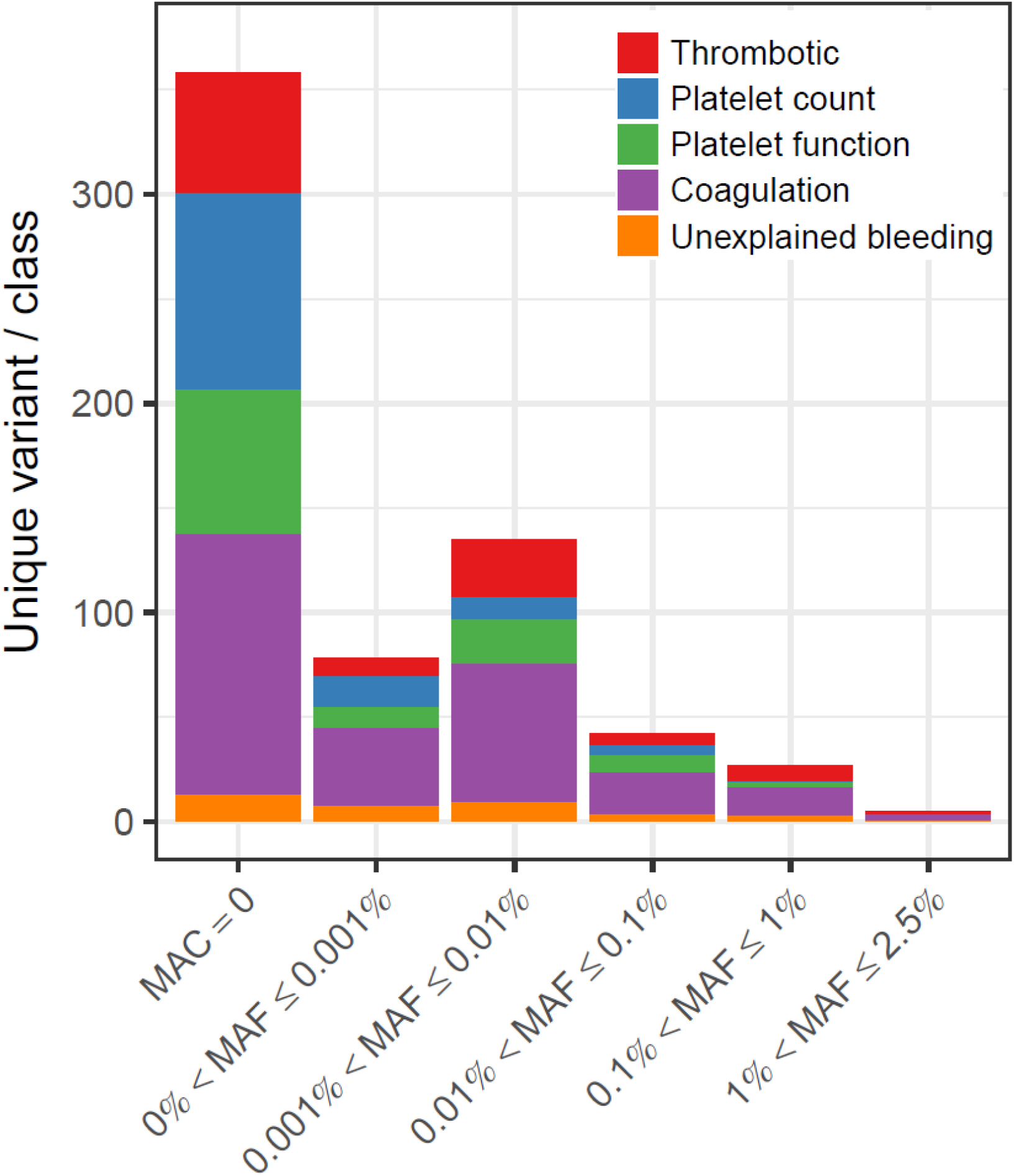
Minor allele frequency or allele count (gnomAD) for the reported autosomal SNVs and indels for each disease class. MAC: Minor Allele Count. MAF: Minor Allele Frequency. The first bin in the plot (MAC=0) corresponds to variants not observed in gnomAD.

**Supplemental Figure 5.**
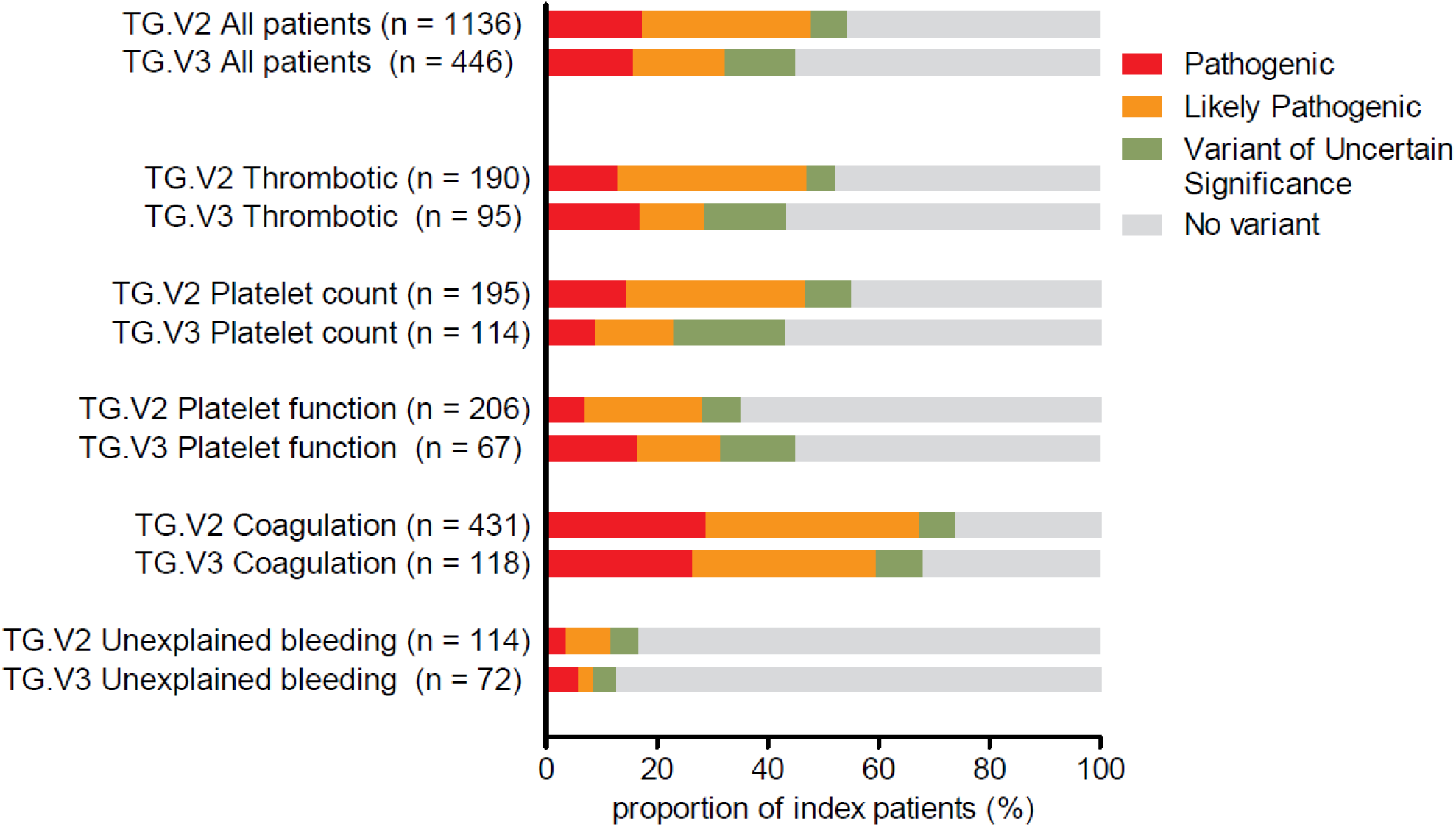
Diagnostic yield and pathogenicity of variants reported using the TG.V2 and TG.V3 test. Only index patients from the ThromboGenomics collection were used for this analysis as both the TG.V2 and TG.V3 platforms were used for to sequence samples from these patients (TG.V2; 1,136 patients and TG.V3; 466 patients).

**Supplemental Figure 6.**
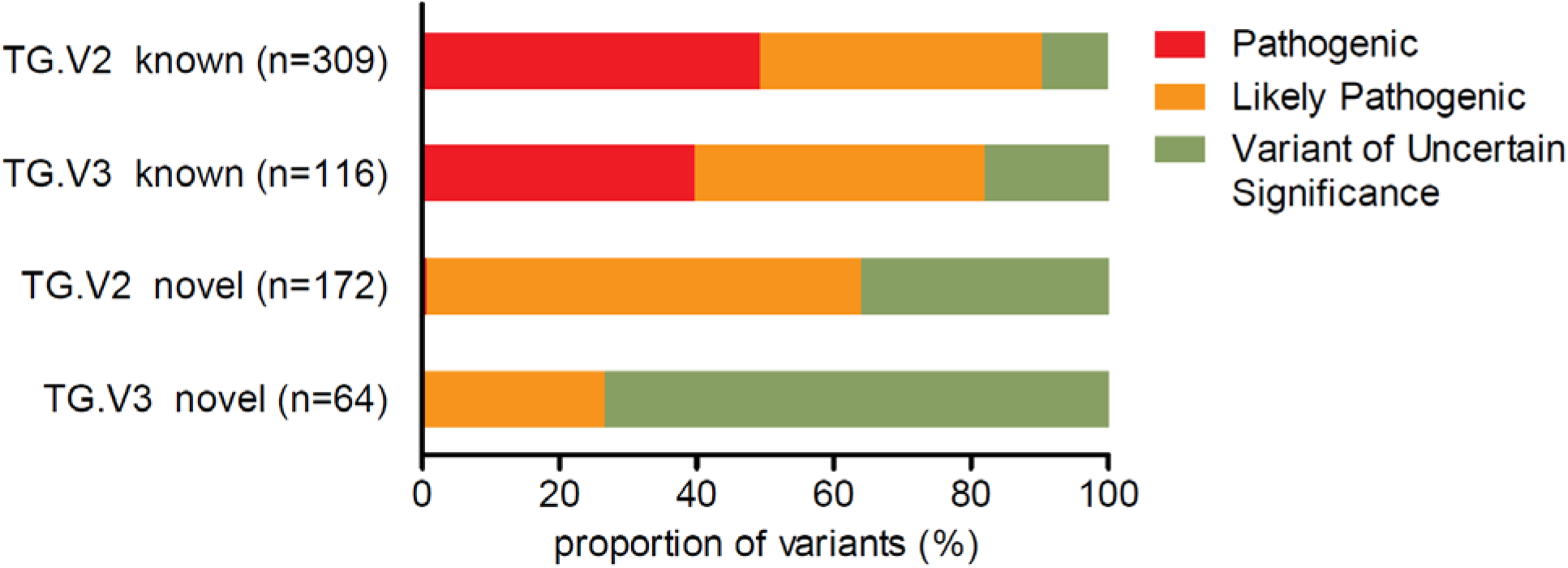
The pathogenicity of novel and known missense variants reported for TG.V2 and TG.V3 in patients from the ThromboGenomics collection. ACMG guidelines were followed for TG.V3 patients. Analysis included 661 missense variants reported in index patients using TG.V2 (481 variants) and TG.V3 (180 variants).

**Supplemental Figure 7.**
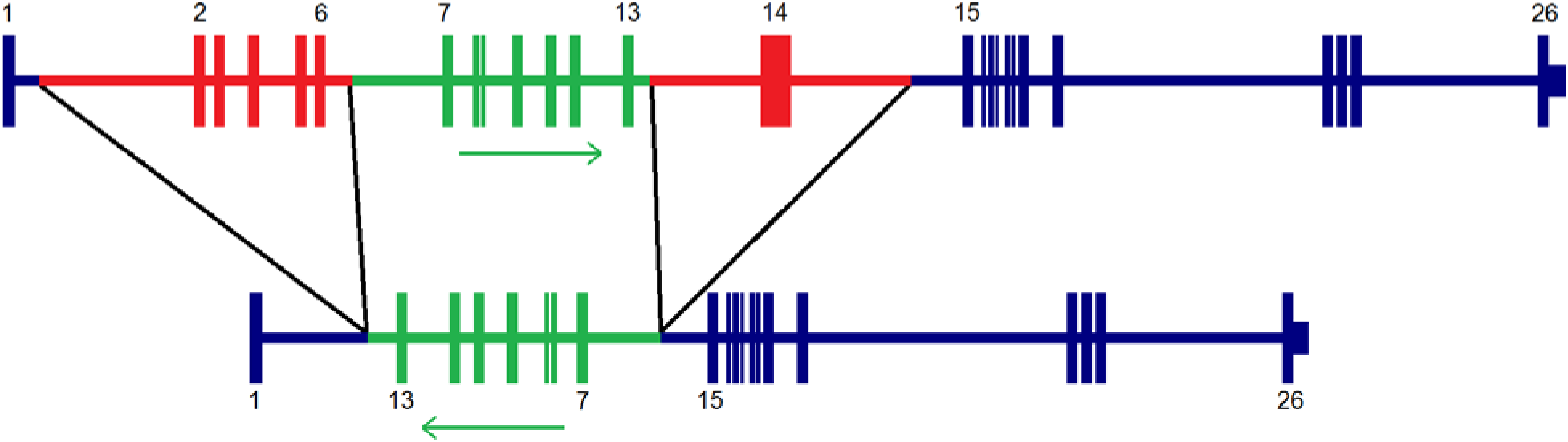
Hemizygous complex CNV associated with severe hemophilia A identified using the ThromboGenomics HTS test. Deletion of intron 1 to intron 6 (red) and intron 13 to intron 14 (red), flanking an inversion of intron 6 to intron 13 (green).

**Supplemental Figure 8.**
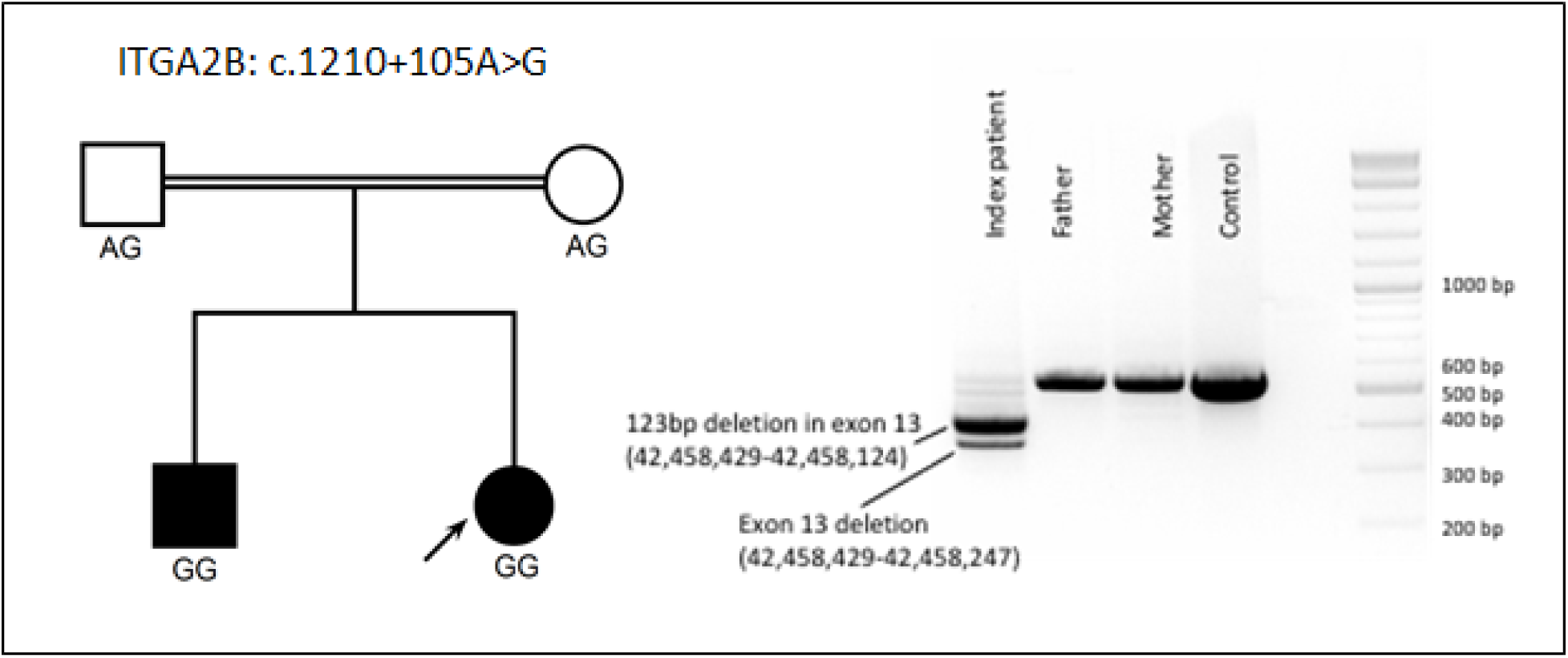
Pedigree of proband with Glanzmann Thrombasthenia caused by a homozygous deep intronic variant. Electrophoresis gel results of PCR amplicons of platelet cDNA reveals that aberrant splicing of the *ITGA2B* mRNA is associated with the deep intronic variant (c.1210+105A>G). Sanger sequencing of the aberrant splice products revealed two mRNAs, one with a deletion of all of exon 13 coding sequence and the second with a 123 bp deletion in the exon 13 coding sequence.

**Supplemental Table 3.**
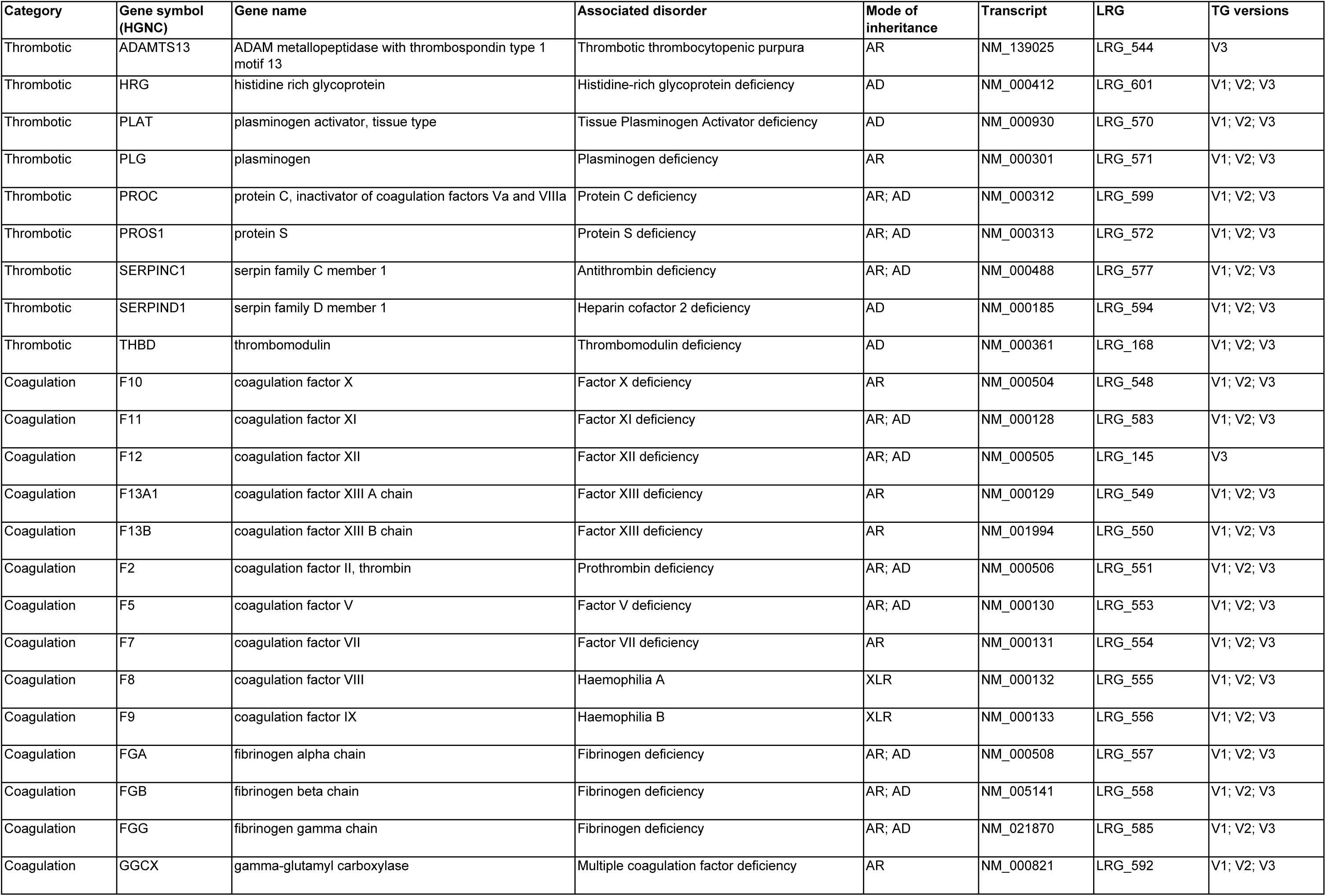
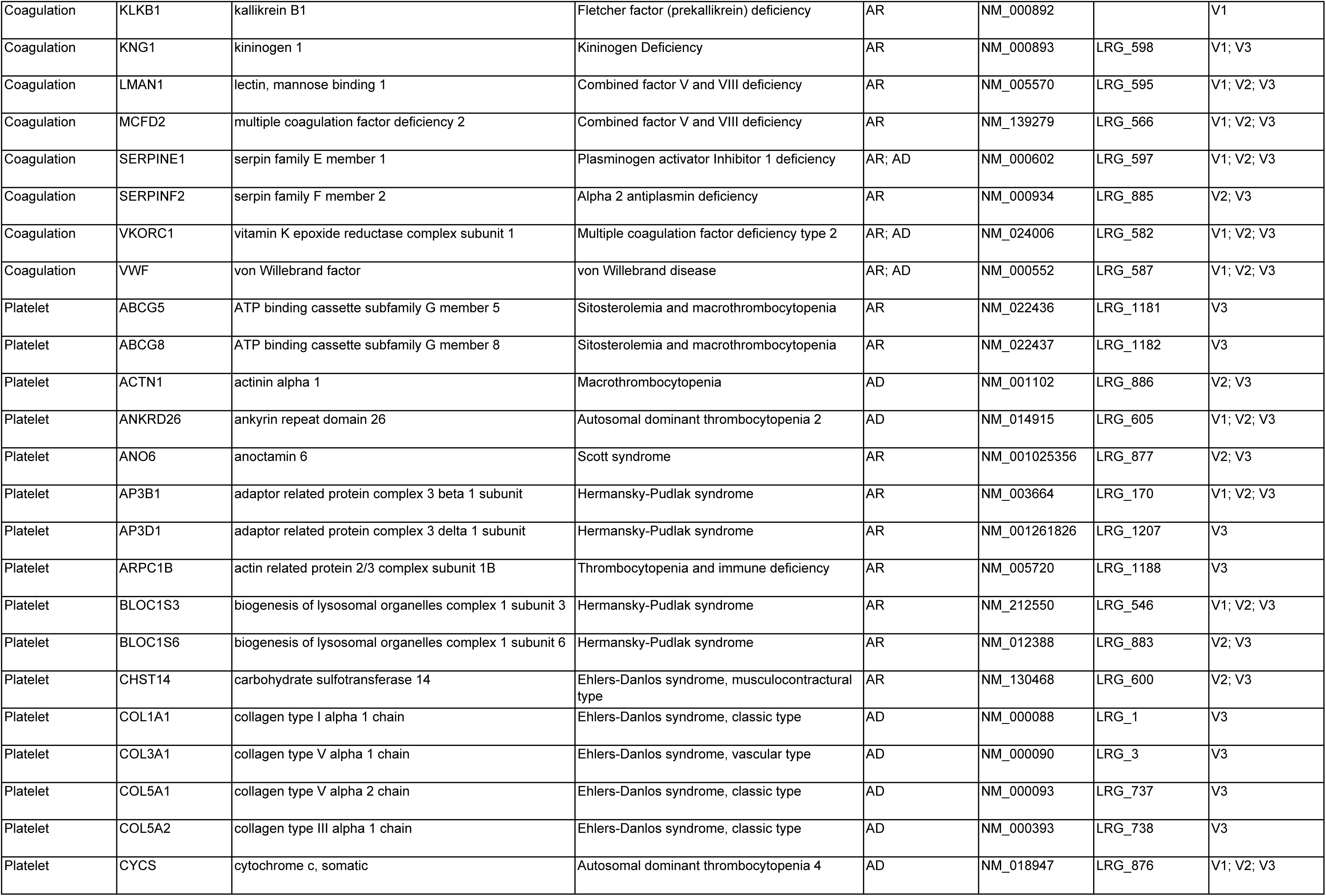
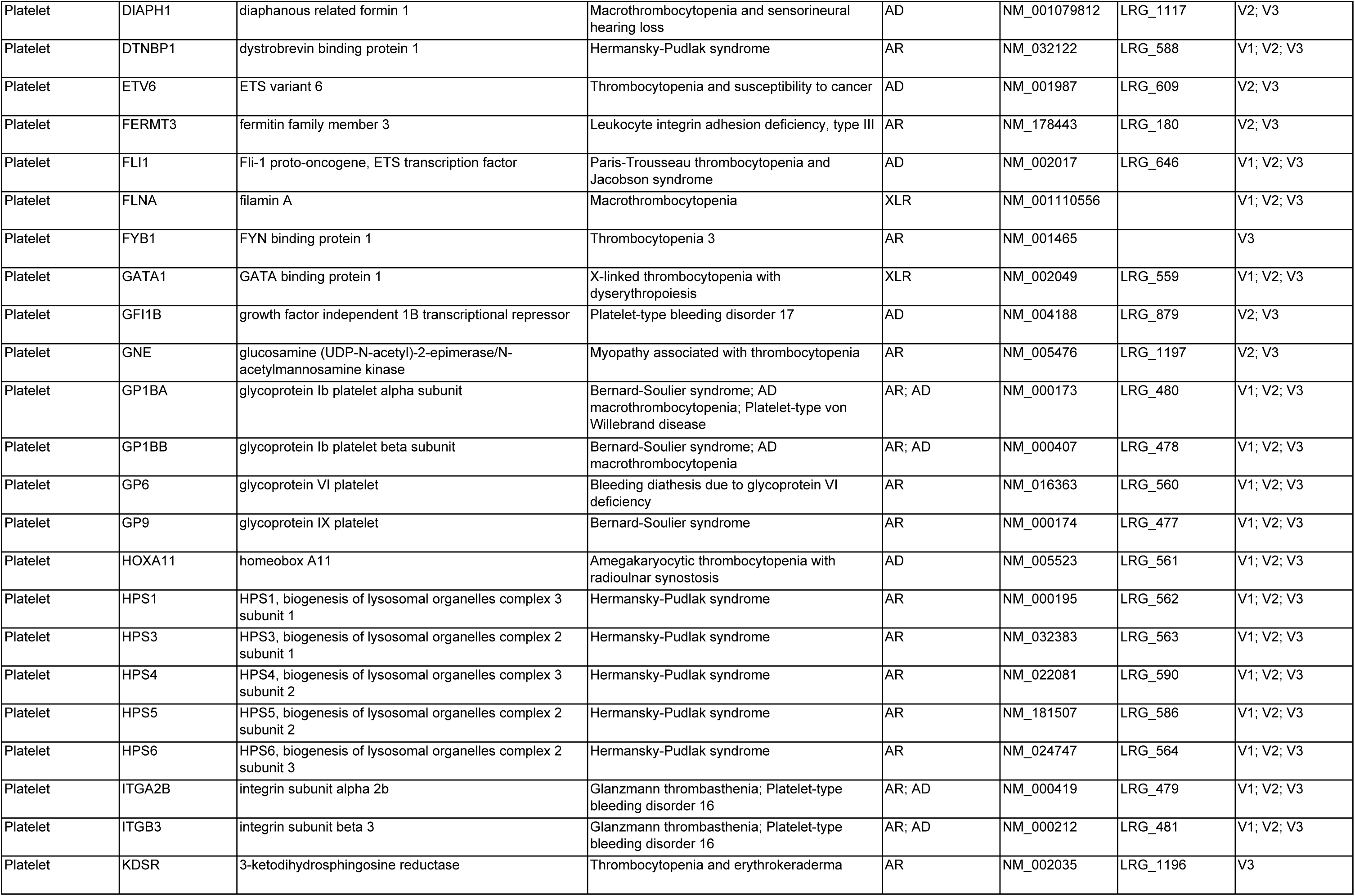
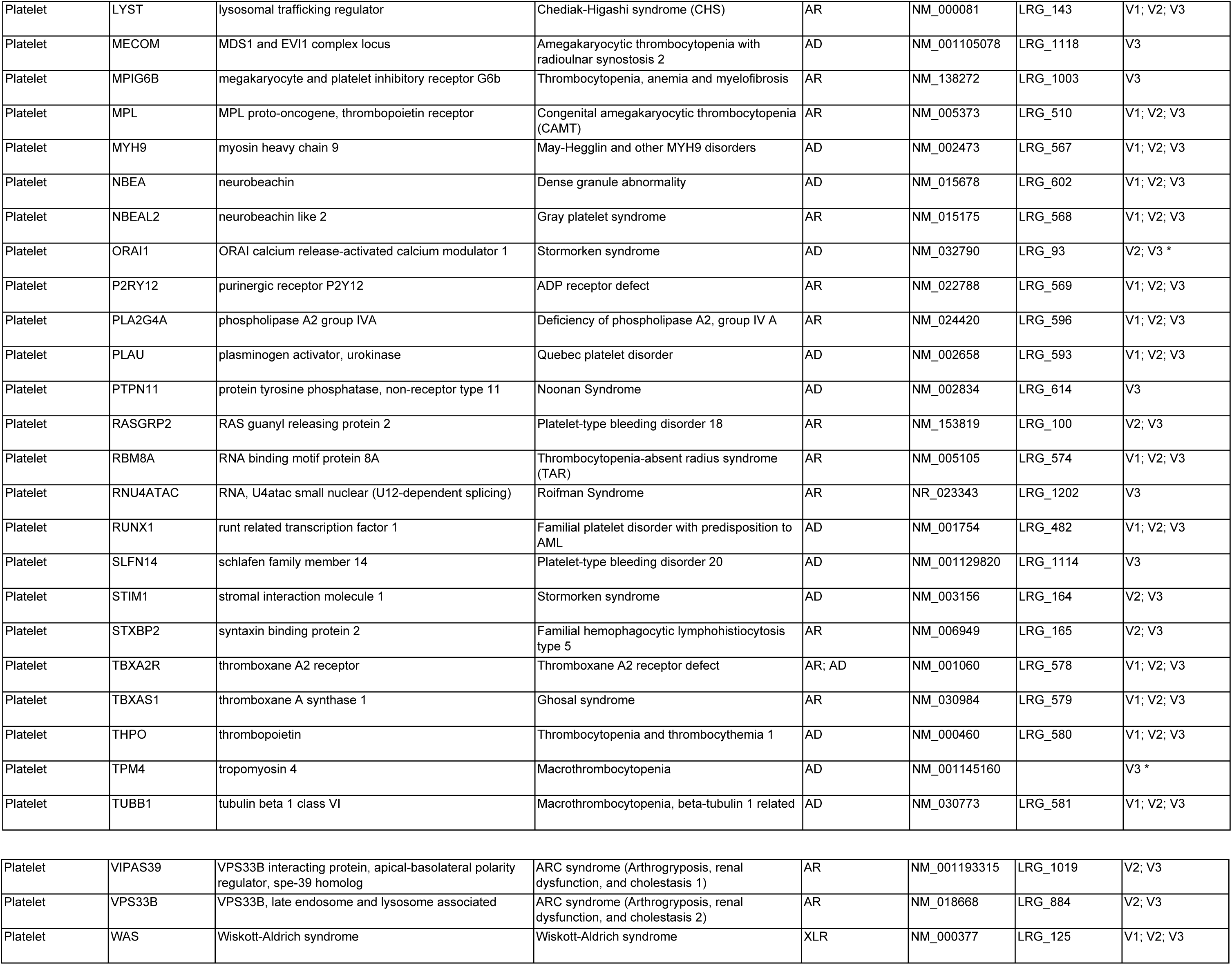

## References

1. Castaman G, Linari S. Diagnosis and Treatment of von Willebrand Disease and Rare Bleeding Disorders. J Clin Med. 2017;6(4).

2. UKHCDO. UKHCDO Annual Report. 2017.

3. Khan S, Dickerman JD. Hereditary thrombophilia. Thromb J. 2006;4:15.

4. Simeoni I, Stephens JC, Hu F, et al. A high-throughput sequencing test for diagnosing inherited bleeding, thrombotic, and platelet disorders. Blood. 2016;127(23):2791–2803.

5. Johnson B, Lowe GC, Futterer J, et al. Whole exome sequencing identifies genetic variants in inherited thrombocytopenia with secondary qualitative function defects. Haematologica. 2016;101(10):1170–1179.

6. Bastida JM, Lozano ML, Benito R, et al. Introducing high-throughput sequencing into mainstream genetic diagnosis practice in inherited platelet disorders. Haematologica. 2018;103(1):148–162.

7. Leinoe E, Zetterberg E, Kinalis S, et al. Application of whole-exome sequencing to direct the specific functional testing and diagnosis of rare inherited bleeding disorders in patients from the Oresund Region, Scandinavia. Br J Haematol. 2017;179(2):308–322.

8. Lee EJ, Dykas DJ, Leavitt AD, et al. Whole-exome sequencing in evaluation of patients with venous thromboembolism. Blood Adv. 2017;1(16):1224–1237.

9. Johnson B, Doak R, Allsup D, et al. A comprehensive targeted next-generation sequencing panel for genetic diagnosis of patients with suspected inherited thrombocytopenia. Res Pract Thromb Haemost. 2018;2(4):640–652.

10. Richards S, Aziz N, Bale S, et al. Standards and guidelines for the interpretation of sequence variants: a joint consensus recommendation of the American College of Medical Genetics and Genomics and the Association for Molecular Pathology. Genet Med. 2015;17(5):405–424.

11. Kohler S, Vasilevsky NA, Engelstad M, et al. The Human Phenotype Ontology in 2017. Nucleic Acids Res. 2017;45(D1):D865–D876.

12. Westbury SK, Turro E, Greene D, et al. Human phenotype ontology annotation and cluster analysis to unravel genetic defects in 707 cases with unexplained bleeding and platelet disorders. Genome Med. 2015;7(1):36.

13. Vries MJ, van der Meijden PE, Kuiper GJ, et al. Preoperative screening for bleeding disorders: A comprehensive laboratory assessment of clinical practice. Res Pract Thromb Haemost. 2018;2(4):767–777.

14. Moenen F, Vries MJA, Nelemans PJ, et al. Screening for platelet function disorders with Multiplate and platelet function analyzer. Platelets. 2017:1–7.

15. Gebhart J, Hofer S, Panzer S, et al. High proportion of patients with bleeding of unknown cause in persons with a mild-to-moderate bleeding tendency: Results from the Vienna Bleeding Biobank (VIBB). Haemophilia. 2018;24(3):405–413.

16. Stenson PD, Mort M, Ball EV, et al. The Human Gene Mutation Database: towards a comprehensive repository of inherited mutation data for medical research, genetic diagnosis and next-generation sequencing studies. Hum Genet. 2017;136(6):665–677.

17. Lek M, Karczewski KJ, Minikel EV, et al. Analysis of protein-coding genetic variation in 60,706 humans. Nature. 2016;536(7616):285–291.

18. Noris P, Pecci A. Hereditary thrombocytopenias: a growing list of disorders. Hematology Am Soc Hematol Educ Program. 2017;2017(1):385–399.

19. Simeoni I, Shamardina O, Deevi SV, et al. GRID - Genomics of Rare Immune Disorders: a highly sensitive and specific diagnostic gene panel for patients with primary immunodeficiencies. bioRxiv. 2018.

20. Landrum MJ, Lee JM, Benson M, et al. ClinVar: improving access to variant interpretations and supporting evidence. Nucleic Acids Res. 2018;46(D1):D1062–D1067.

21. Westbury SK, Downes K, Burney C, et al. Phenotype description and response to thrombopoietin receptor agonist in DIAPH1-related disorder. Blood Adv. 2018;2(18):2341–2346.

22. Albers CA, Paul DS, Schulze H, et al. Compound inheritance of a low-frequency regulatory SNP and a rare null mutation in exon-junction complex subunit RBM8A causes TAR syndrome. Nat Genet. 2012;44(4):435–439, S431–432.

23. England G. The 100,000 Genomes Project Protocol. v.3. 2017(version3).

24. Poggi M, Canault M, Favier M, et al. Germline variants in ETV6 underlie reduced platelet formation, platelet dysfunction and increased levels of circulating CD34+ progenitors. Haematologica. 2017; 102(2):282–294.

25. Chen L, Kostadima M, Martens JHA, et al. Transcriptional diversity during lineage commitment of human blood progenitors. Science. 2014;345(6204):1251033.

26. Revel-Vilk S, Shai E, Turro E, et al. GNE variants causing autosomal recessive macrothrombocytopenia without associated muscle wasting. Blood. 2018;132(17):1851–1854.

27. Futterer J, Dalby A, Lowe GC, et al. Mutation in GNE is associated with severe congenital thrombocytopenia. Blood. 2018;132(17):1855–1858.

28. Westbury SK, Canault M, Greene D, et al. Expanded repertoire of RASGRP2 variants responsible for platelet dysfunction and severe bleeding. Blood. 2017;130(8):1026–1030.

29. Sivapalaratnam S, Westbury SK, Stephens JC, et al. Rare variants in GP1BB are responsible for autosomal dominant macrothrombocytopenia. Blood. 2017;129(4):520–524.

30. Inaba H, Shinozawa K, Amano K, Fukutake K. Identification of deep intronic individual variants in patients with hemophilia A by next-generation sequencing of the whole factor VIII gene. Res Pract Thromb Haemost. 2017;1(2):264–274.

31. Nuzzo F, Bulato C, Nielsen BI, et al. Characterization of an apparently synonymous F5 mutation causing aberrant splicing and factor V deficiency. Haemophilia. 2015;21(2):241–248.

32. Daidone V, Gallinaro L, Grazia Cattini M, et al. An apparently silent nucleotide substitution (c.7056C>T) in the von Willebrand factor gene is responsible for type 1 von Willebrand disease. Haematologica. 2011;96(6):881–887.

33. Xie J, Pabon D, Jayo A, Butta N, Gonzalez-Manchon C. Type I Glanzmann thrombasthenia caused by an apparently silent beta3 mutation that results in aberrant splicing and reduced beta3 mRNA. Thromb Haemost. 2005;93(5):897–903.

34. Pippucci T, Savoia A, Perrotta S, et al. Mutations in the 5’ UTR of ANKRD26, the ankirin repeat domain 26 gene, cause an autosomal-dominant form of inherited thrombocytopenia, THC2. Am J Hum Genet. 2011;88(1):115–120.

35. Sabater-Lleal M, Chillon M, Howard TE, et al. Functional analysis of the genetic variability in the F7 gene promoter. Atherosclerosis. 2007;195(2):262–268.

36. Petersen R, Lambourne JJ, Javierre BM, et al. Platelet function is modified by common sequence variation in megakaryocyte super enhancers. Nat Commun. 2017;8:16058.

37. Rehm HL, Berg JS, Brooks LD, et al. ClinGen-the Clinical Genome Resource. N Engl J Med. 2015;372(23):2235–2242.

38. Bycroft C, Freeman C, Petkova D, et al. The UK Biobank resource with deep phenotyping and genomic data. Nature. 2018;562(7726):203–209.

39. Turnbull C, Scott RH, Thomas E, et al. The 100 000 Genomes Project: bringing whole genome sequencing to the NHS. BMJ. 2018;361:k1687.

40. Abraham G, Havulinna AS, Bhalala OG, et al. Genomic prediction of coronary heart disease. Eur Heart J. 2016;37(43):3267–3278.

41. Khera AV, Emdin CA, Drake I, et al. Genetic Risk, Adherence to a Healthy Lifestyle, and Coronary Disease. N Engl J Med. 2016;375(24):2349–2358.

42. Fuchsberger C, Flannick J, Teslovich TM, et al. The genetic architecture of type 2 diabetes. Nature. 2016;536(7614):41–47.

43. Maas P, Barrdahl M, Joshi AD, et al. Breast Cancer Risk From Modifiable and Nonmodifiable Risk Factors Among White Women in the United States. JAMA Oncol. 2016;2(10):1295–1302.

## Supplemental References

1. Simeoni I, Stephens JC, Hu F, et al. A high-throughput sequencing test for diagnosing inherited bleeding, thrombotic, and platelet disorders. Blood. 2016;127(23):2791–2803.

2. Rossetti LC, Radic CP, Larripa IB, De Brasi CD. Developing a new generation of tests for genotyping hemophilia-causative rearrangements involving int22h and int1h hotspots in the factor VIII gene. J Thromb Haemost. 2008;6(5):830–836.

3. Vries MJ, van der Meijden PE, Kuiper GJ, et al. Preoperative screening for bleeding disorders: A comprehensive laboratory assessment of clinical practice. Res Pract Thromb Haemost. 2018;2(4):767–777.

4. Moenen F, Vries MJA, Nelemans PJ, et al. Screening for platelet function disorders with Multiplate and platelet function analyzer. Platelets. 2017:1–7.

5. Gebhart J, Hofer S, Panzer S, et al. High proportion of patients with bleeding of unknown cause in persons with a mild-to-moderate bleeding tendency: Results from the Vienna Bleeding Biobank (VIBB). Haemophilia. 2018;24(3):405–413.

6. Stenson PD, Mort M, Ball EV, et al. The Human Gene Mutation Database: towards a comprehensive repository of inherited mutation data for medical research, genetic diagnosis and next-generation sequencing studies. Hum Genet. 2017;136(6):665–677.

7. Fokkema IF, Taschner PE, Schaafsma GC, Celli J, Laros JF, den Dunnen JT. LOVD v.2.0: the next generation in gene variant databases. Hum Mutat. 2011;32(5):557–563.

8. McKenna A, Hanna M, Banks E, et al. The Genome Analysis Toolkit: a MapReduce framework for analyzing next-generation DNA sequencing data. Genome Res. 2010;20(9):1297–1303.

9. Cingolani P, Platts A, Wang le L, et al. A program for annotating and predicting the effects of single nucleotide polymorphisms, SnpEff: SNPs in the genome of Drosophila melanogaster strain w1118; iso-2; iso-3. Fly (Austin). 2012;6(2):80–92.

10. Plagnol V, Curtis J, Epstein M, et al. A robust model for read count data in exome sequencing experiments and implications for copy number variant calling. Bioinformatics. 2012;28(21):2747–2754.

